# Symbiotic versatility in action: *Trebouxia* diversity expands the niche of the lichen *Xanthoria parietina*

**DOI:** 10.1101/2025.11.12.687866

**Authors:** Salvador Chiva, Tamara Pazos, Javier Montero-Pau, Patricia Moya, Isaac Garrido-Benavent, Eva Barreno, Lucia Muggia

## Abstract

**Aim:** We explored the phylogenetic diversity of the mycobiont *Xanthoria parietina* and its associated *Trebouxia* phycobionts across the Mediterranean region and examined how photobiont climatic preferences may shape the ecological amplitude and spatial distribution of the lichen.

**Location:** Specimens were collected in different localities in the central and western Mediterranean Basin (Iberian and Italian peninsulas), with additional Atlantic and boreal localities used for comparison.

**Methods:** Genetic diversity and phylogenies of both lichen symbionts were inferred from nrITS data. Mycobiont–phycobiont interaction networks were constructed, and ecological niches of associated *Trebouxia* species were modelled using 19 bioclimatic variables from WorldClim v. 2.1.

**Results:** Phylogenetic analyses revealed high diversity within *Xanthoria parietina* and clarified the placement of poorly studied species within the genus *Xanthoria* (e.g., *X. monofoliosa* and *X. aureola* s. lat.), and revealed a novel lineage (*Xanthoria* sp. ‘hydra’). All the photobionts belonged to *Trebouxia* clade A; nine *Trebouxia* species-level lineages were detected, including a previously unrecognized lineage (*Trebouxia* sp. A56). *Trebouxia decolorans* (A33) was the most frequent photobiont and exhibited the broadest climatic niche, whereas *T. solaris* (A35) and *T. tabarcae ad int.* (A48) showed a more specialized climatic niche. Ensemble species-distribution models indicated widespread suitability for *T. decolorans* across Europe, and more coastal-Mediterranean suitability for *T. tabarcae ad int*.

**Main Conclusion:** *Xanthoria parietina* displays remarkable symbiont flexibility in associating with various *Trebouxia* lineages within clade A, which contribute to its ecological success in heterogeneous Mediterranean environments. Climatic niche modelling suggests that the association with different phycobionts may expand the ecological niche of the lichen, potentially enhancing its capacity to inhabit a wider range of environmental conditions.

## 1. Introduction

Lichens are multidimensional, long-lived symbiotic systems that integrate biotic, morphological, and functional entities (Margulis 1993). Lichen thalli emerge as ’holobiomes’ because they result from specific interactions and the subsequent integration of a diverse array of microorganisms (Margulis and Barreno 2003). These thalli involve, as key players, a principal heterotrophic fungus, the mycobiont, which provides a stable structure, along with main unicellular photosynthetic partners (the photobionts), which can be green microalgae (phycobionts) and/or cyanobacteria (Honegger 1998). Additionally, other microorganisms, such as non-photosynthetic bacteria, lichenicolous fungi (either filamentous or yeasts), and other microalgae, have been identified as integral components of the lichen symbiosis (Casano et al. 2011; Cometto et al. 2023; Hawksworth and Grube 2020; Moya et al. 2017; Spribille 2018).

Among the phycobionts, green microalgae of the genus *Trebouxia* Puymaly play a prominent role, as they associate with 80% of the known lichen-forming fungi in temperate regions and over 20% globally (Friedl and Büdel 2008; Friedl and Gärtner 1988; Hawksworth and Lücking 2017; Rambold et al. 1998). *Trebouxia* represents one of the major lineages within the class *Trebouxiophyceae* (Muggia et al. 2018). However, despite the formal taxonomic recognition of 34 species within the genus, many species-level lineages remain undescribed due to the difficulties associated with isolating and culturing these microalgae outside their symbiotic state (Chiva et al. 2025; Muggia et al. 2018, 2020; Pazos et al. 2025; Veselá et al. 2024). To address this issue, an integrative taxonomic approach incorporating genetic, physiological, and morphological data has been adopted (Barreno et al. 2022; Pazos et al. 2025), utilizing the coding system established by Leavitt et al. (2015) and Muggia et al. (2020) for species-level lineages of *Trebouxia*. Within this system, *Trebouxia* lineages are classified into four main clades (A, C, I, and S; Beck 2002; Muggia et al. 2020), while a fifth clade (clade D) has only recently been established to accommodate a group identified exclusively through sequence data (Xu et al. 2020). According to the *Trebouxia* Research Portal (https://trebouxia.net/; last update July 2025), which compiles representative sequences of each *Trebouxia* lineage, clade A is the most diverse, encompassing over 70 lineages, followed by clade C with 45, clade I with 32, clade S with 28, and clade D with six.

Understanding the ecological niche characteristics of the *Trebouxia* lineages is crucial to assess their adaptive strategies within lichen symbioses (Nelsen et al. 2021, 2022). Recent studies have revealed that photobiont variability influences lichen adaptation to different climatic conditions, leading to significant niche expansions, as observed in associations with *Lasallia* spp. (Rolshausen et al. 2018) and *Ramalina farinacea* (Moya et al. 2024). Climatic niche modelling approaches have shown that different *Trebouxia* species exhibit varying levels of specialization and generalization (Kosecka et al. 2022; Nelsen et al. 2021, 2022; Pazos et al. 2025). However, the number of *Trebouxia* species for which climatic niche modelling has been applied remains very limited. Even abundant and widespread lineages, such as *T. decolorans*, have never been evaluated using these approaches. Significant progress in this field has been achieved through studies on *Asterochloris* species, where ecological niche modelling has revealed patterns of niche differentiation and partitioning among closely related lineages (e.g., Vančurová et al. 2018, 2021). These studies highlight the potential of niche modelling to uncover ecological strategies in photobionts. In this context, climatic niche modelling suggests that the association with different phycobionts may expand the ecological niche of the lichen, enhancing its adaptability to diverse environments.

An outstanding example of a highly versatile and cosmopolitan lichen is *Xanthoria parietina* (L.) Ach. (*Teloschistaceae*), characterized by its bright orange-yellow foliose thallus. This species, globally distributed across coastal and mountainous regions, is remarkably resilient to environmental stress and has even been designated as a “Martian lichen” due to its potential adaptability to Mars-like conditions—demonstrated by its ability to survive and remain metabolically active after exposure to high levels of UV and cosmic radiation (Lorenz et al. 2023, 2024). It thrives on nitrogen-rich substrates, including nutrient-rich limestones and siliceous rocks, walls, fences, roofs, and high-pH tree barks (Nimis et al. 2025; Silberstein et al. 1996). Geographically, *X. parietina* has been reported in Europe, Asia, Australia, Africa, and North and South America (Aptroot 2008; Calvelo and Liberatore 2002; Galloway and Quilhot 1998; Lindblom and Ekman 2005; Malme 1926). Its broad ecological amplitude and global success make it a valuable model for studying how symbiotic association with different phycobionts can influence niche breadth and geographic distribution in lichens.

Furthermore, *Xanthoria parietina* primarily associates with the most common *Trebouxia* species, such as *T. aggregata*, *T. arboricola*, *T. crenulata*, and *T. decolorans* (Beck et al. 1998; Beck and Mayr 2012; Dal Grande et al. 2014; Nyati et al. 2013, 2014; Voytsekhovich and Beck 2016; Wyczanska et al. 2023). Some authors observed that saxicolous or epilithic thalli of *X. parietina* are associated with *T. arboricola*, whereas *T. decolorans* is the preferred phycobiont in epiphytic thalli (Nyati et al. 2013, 2014). However, Beck and Mayr (2012) did not find any correlation between *X. parietina* lineages, *T. decolorans* microalgae, substrate preferences, isotopic composition, or geographic origin.

*Xanthoria parietina* also stands out for its high resistance and tolerance to abiotic stresses such as salinity, heavy metals, drought and extreme temperatures (Benhamada et al. 2023; Dzubaj et al. 2008; Grimm et al. 2021; Honegger 2001; Lorenz et al. 2023). This resistance allows the lichen to thrive under multiple environmental conditions (known as ecological adaptability, Bertazzo-Silva et al. 2025) and to spread across diverse biogeographic regions (Honegger et al. 2004, Lindblom and Ekman 2005, 2006). Furthermore, *X. parietina* has been reported in areas with high levels of ammonia and nitrate pollution (Barreno and Pérez-Ortega 2003; Gasulla et al. 2025; Giordani et al. 2014; Munzi et al. 2012; Olsen et al. 2010) and has thus been chosen as a key species for biomonitoring studies of air pollution (e.g., Loppi et al. 2006; Moya et al. 2023; Niepsch et al. 2021;). Recently, Gasulla et al. (2025) studied various lichens in urban and seminatural areas, including *X. parietina,* and demonstrated that the combination and interaction of various ecophysiological traits, including nitrophily, osmotic stress, desiccation tolerance, and recovery capacity, determine the ability of lichen phycobionts to survive in harsh, and nitrogen-polluted environments.

Despite the fact that about 10,000 articles have been published on *Xanthoria parietina* (https://scholar.google.com/), only a few have focused on its photobionts and the mycobiont-photobiont interaction patterns (Beck and Mayr 2012; Dal Grande et al. 2014; Nyati et al. 2013, 2014; Wyczanska et al. 2023). Furthermore, *X. parietina*, in general, builds thalli that are easily recognizable in the field. However, due to its ability to grow on different substrates and to associate with different *Trebouxia* species, the mycobiont itself may conceal a certain degree of genetic diversity—or even represent a complex of cryptic species, or taxa, that are easily misidentified and commonly grouped under the name *Xanthoria parietina*.

In this study, we aimed to explore the phylogenetic diversity of the mycobiont *Xanthoria parietina*, and the genetic diversity of its associated primary *Trebouxia* species-level lineages. Samples were collected from different localities across the Mediterranean Basin, both in the Iberian and Italian peninsulas. In addition, the climatic preferences of the *Trebouxia* species-level lineages were characterized with the aim of understanding how the association with different algal partners may contribute to expanding the overall ecological niche and potential distribution range of the *X. parietina* symbiosis. It also may predicts the possible effects of climate change.

## 2. Material and Methods

### 2.1. Sample collection

A total of 152 lichen samples, visually identified as *Xanthoria parietina*, were collected from 36 localities in Mediterranean biogeographic and bioclimatic regions across the Iberian Peninsula (IbP) and the Italian Peninsula (ItP). Of these, 72 samples were collected from 17 separate localities in the IbP, while 80 samples were collected from 19 localities in the ItP. Additionally, 22 thalli of *X. parietina* were collected from other European biogeographic and bioclimatic areas (Figure 1, Table S1): in the Atlantic (two localities in France and two in the northern Iberian Peninsula—one in Navarra and one in Euskadi), in Central European (one in Slovakia) and in the Boreal (one in Norway). All the collected samples were assigned an alphanumeric collection code corresponding to the first author, abbreviated as SCH. The biogeographic regions of the sampling localities were recognized according to Rivas-Martínez et al. (2017).

Only ten samples were saxicolous thalli (growing on rocks), while the rest were epiphytic, growing on various phorophyte species (Table S1). Nevertheless, these were also included in the study.

**FIGURE 1.**
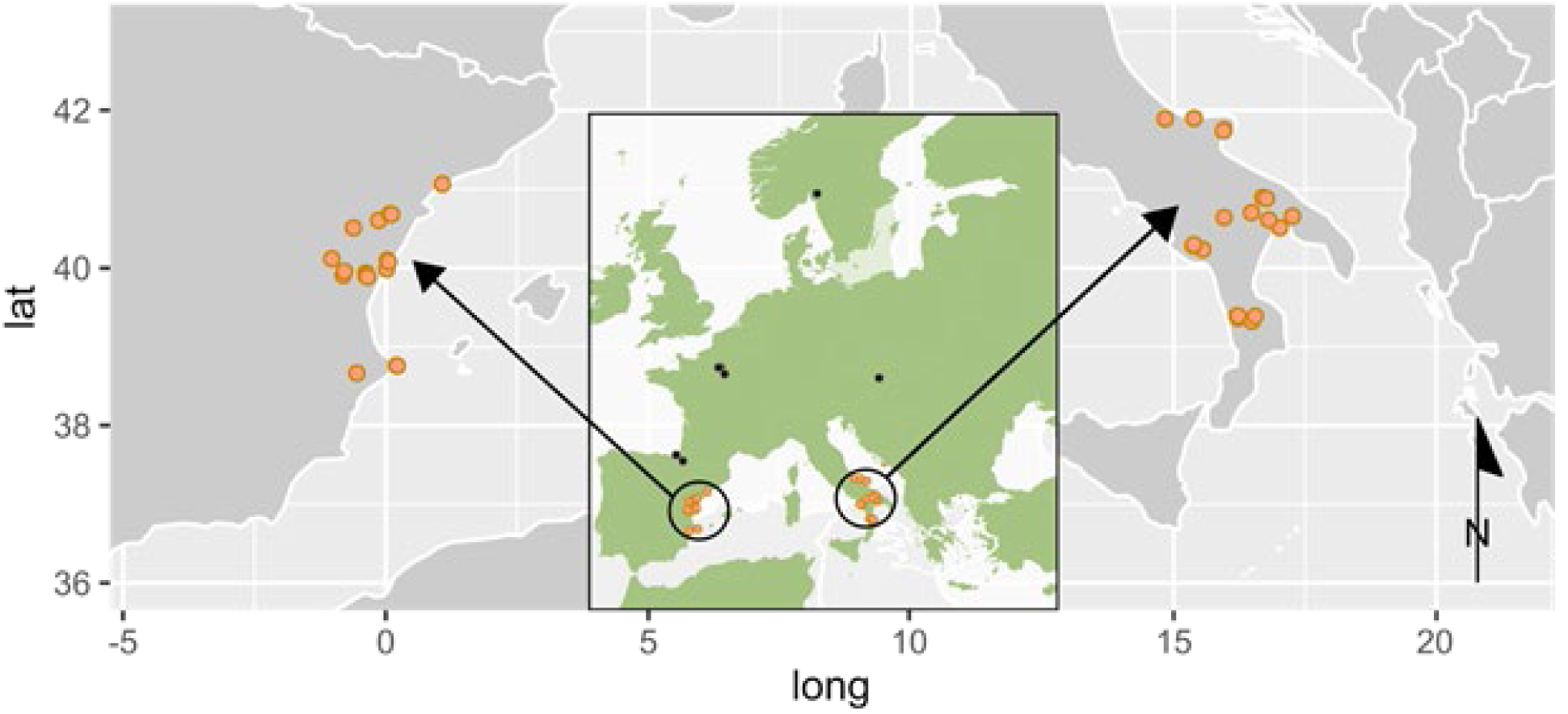
Sampling localities of *Xanthoria parietina* thalli across different biogeographic regions. The two areas enclosed within a circle correspond to the Mediterranean sampling sites (indicated by orange dots), located in the Iberian and Italian Peninsulas. Localities from other biogeographic regions in central and northern Europe are indicated by black dots.

### 2.2. Pretreatment of the samples and DNA extraction

Thalli of *Xanthoria parietina* were screened for the absence of infection symptoms by lichenicolous fungi, and a piece of each thallus was taken for DNA extraction and stored in a 1.5 ml tube. To eliminate possible external contamination, the fragments were washed with 500 μl of distilled water added to each Eppendorf tube, agitated for 5 minutes, and then the supernatant was discarded. Subsequently, 500 μl of a solution of distilled water and Tween20 was added, agitated for 30 minutes, after which two washes with distilled water were performed by briefly vortexing. The lichen fragments were then transferred into a new tube and DNA extraction followed the protocol of Cubero et al. (1999) adapted for lichens, i.e., the extraction began by freezing and pulverizing the samples with liquid nitrogen and micropestles.

### 2.3. PCR amplification and sequencing

PCR amplification was performed using specific primers for the internal transcribed spacer of the nuclear ribosomal DNA (nrITS) of both the mycobiont and the phycobiont. Initially, the mycobiont primers ITS1F (forward, Gardes and Bruns 1993) and ITS4A (reverse, Larena et al. 1999) were used with a subset of samples to verify their effectiveness. The results showed that less than 50% of the sequences were usable, prompting the design of a forward primer specific to the mycobiont *X. parietina*, named Xantho-F (5’-GATCATTACCGAGAGTGACGGA-3’). The newly designed Xantho-F primer, together with ITS4A, was then used. For the microalgae, the selected primer pair was nr-SSU-1780-5’ (Piercey-Normore and Depriest 2001) and ITS4T (Kroken and Taylor 2000).

PCR amplifications began with an initial denaturation step of 2 min at 94 °C, followed by denaturation for 30 sec at 94 °C, primer annealing for 45 sec at 58 °C, and extension at 72 °C for 1 min. This process was repeated for 30 cycles and ended with a final extension step at 72 °C for 7 min. For both primer pairs, PCR amplifications were carried out in a total volume of 25 μL, containing 12.5 μL of EmeraldAmp GT PCR Master Mix (Takara), 0.5 μL of each primer (10 μM), 1 μL of dimethyl sulfoxide (DMSO), 1 μL of DNA, and 9.5 μL of sterile Milli-Q water. PCR products were visualized on 1% agarose gels and purified using the NZYGelPure kit (NZYTech) following the manufacturer’s instructions. Purified PCR products were sequenced using an ABI 3100 genetic analyzer with the ABI BigDyeTM Terminator cycle sequencing kit (Applied Biosystems, Foster City, California) at StabVida (Portugal).

### 2.4. Phylogenetic analyses and interaction networks

Preliminary taxonomic identification to the genus level of myco- and phycobionts was performed by BLAST search of all the nrITS sequences obtained against the GenBank nucleotide database (Altschul et al. 1990; https://www.ncbi.nlm.nih.gov/genbank/). Subsequently, a phylogenetic analysis of the *Xanthoria* mycobionts was conducted, for which a multiple sequence alignment (MSA) was first built including the 174 newly obtained nrITS sequences, along with a selection of 78 *Xanthoria* sequences retrieved from the GenBank, most of which were previously generated in the studies of Beck and Mayr (2012), Khodosovtsev et al. (2023), Scherrer and Honegger (2003) and Tsurykau et al. (2020). A sequence of the species *Martinjahnsia resendei* (AF101285) was chosen as the outgroup to root the phylogeny, as this lichenized fungus belongs to a genus closely related to *Xanthoria* (Kondratyuk et al. 2024).

Because all the *Trebouxia* phycobiont sequences were identified as *Trebouxia* species belonging to clade A, we then built a MSA including the 145 newly obtained nrITS sequences and a selection of 96 sequences retrieved from the GenBank and from the *Trebouxia* Research Portal (https://trebouxia.net/; last update July 2025). These sequences corresponded to the microalgal lineages described in the studies of Barreno et al. (2022), Chiva et al. (2025), Kosecka et al. (2022) and Muggia et al. (2020). To cover the entire *Trebouxia* diversity associated with the genus *Xanthoria*, 131 nrITS sequences of *Trebouxia* deposited in the GenBank as symbionts of *Xanthoria* spp., were also included in the MSA. Many of these sequences were not accurately identified, as they were submitted before the current coding system for *Trebouxia* lineages introduced by Leavitt et al. (2015) and Muggia et al. (2020). These sequences were retrieved from the GenBank using keyword searches (e.g., ’Trebouxia AND Xanthoria AND internal transcribed spacer’). Two sequences of *Trebouxia simplex* (KJ623934 and KJ623927) from *Trebouxia* clade S were used as the outgroup to root the resulting phylogeny.

Multiple sequence alignments were obtained using MAFFT v. 7.505 (Katoh et al. 2002; Katoh and Standley 2013), implementing the FFT-NS-I x1000 algorithm, the 200PAM / k = 2 scoring matrix, a gap open penalty of 1.5 and an offset value of 0.123. Phylogenetic trees were constructed using the Maximum Likelihood (ML) method with RAxML v. 8.2.12 (Stamatakis 2006; Stamatakis et al. 2008) as implemented in the CIPRES v. 3.3 portal (Miller et al. 2010), using the GTRGAMMA as the nucleotide substitution model. One thousand rapid bootstrap pseudoreplicates were conducted to evaluate nodal support, and tree nodes with bootstrap support (BS) values higher than 70% were regarded as significantly supported. Additionally, Bayesian inference (BI) analysis was performed using MrBayes v. 3.2.7 (Ronquist et al. 2012) with two parallel, simultaneous, four-chain runs executed over 5×10^7^ generations starting with a random tree, and sampling after every 500th step. The first 25% of data was discarded as burn-in, and the 50% majority-rule consensus tree and corresponding posterior probabilities (PP) were calculated from the remaining trees. Chain convergence was assessed to ensure that the potential scale reduction factor (PSRF) approached 1.00 and that the average standard deviation of split frequencies (ASDSF) values fell below 0.005.

Nodes with PP equal to or higher than 0.95 were considered as significantly supported. Evolutionary models were calculated using the jModelTest2 program (K80+G and SYM+G for myco- and phycobiont datasets, respectively). The resulting phylogenetic trees were visualized using FigTree (http://tree.bio.ed.ac.uk/software/figtree/), and InkScape (https://inkscape.org/) was used for artwork.

Mycobiont-phycobiont bipartite interaction networks were built with data of 145 lichen thalli for which nrITS sequences for both symbionts were successfully obtained using the function *plotweb* in the R package *bipartite* (Dormann et al. 2008; R Core Team 2024). Three bipartite networks were constructed: mycobiont *vs.* phycobiont; mycobiont *vs.* region (ItP or IbP) and phycobiont *vs.* region (ItP or IbP). The latter two networks were combined into a single figure using InkScape (https://inkscape.org/).

### 2.5. Climatic niche hypervolumes

The climatic niche and environmental preferences of different *Trebouxia* species-level lineages involved in symbiosis with *Xanthoria parietina* were evaluated based on the 19 climatic variables from the WorldClim v. 2.1 database at 2.5 arc-minutes spatial resolution of (Fick and Hijmans 2017). For this analysis, the dataset included only *Trebouxia* lineages identified in at least five different sampling localities, a threshold set to ensure a minimally robust representation and basic statistical reliability in subsequent analyses (Rolshausen et al. 2018). Therefore, analyses considered only the phycobionts *Trebouxia decolorans* (A33), *T. solaris* (A35) and *T. tabarcae ad int.* (A48, Moya et al. submitted). The dataset included a total of 135 *Trebouxia* sequences associated with *X. parietina*, comprising 106 newly obtained from our samples—whose mycobiont was identified as *X. parietina* via nrITS analysis—and 29 additional sequences of *Trebouxia decolorans* (A33) retrieved from the GenBank (see Table S2, in green).

To detect and visualize the ordination of data in the hyperspace, a Principal Component Analysis (PCA) was performed using the function *prcomp* from the package *stats* in R. PCA helps to identify the most important axes of variation in the data, allowing us to understand the relationships between climatic variables and microalgal species, and thus to highlight key environmental gradients that affect species’ occurrences. Differences in the climatic niche of the different *Trebouxia* spp. were also visualized using a non-metric multidimensional scaling (NMDS) analysis. The ordination plot was built using the function *metaMDS* from the R package *vegan* (Oksanen et al. 2008), based on pairwise dissimilarities calculated with the Bray-Curtis distance.

Additionally, niches were represented as n-dimensional hypervolumes based on the 19 climatic variables, along with distance to the sea (m) and altitude (meters above sea level; m a.s.l.), following the Hutchinsonian niche concept (Hutchinson 1957). Climatic hypervolumes were constructed using the multivariate kernel density estimation (Blonder et al. 2014) with the *hypervolume* R package (Karger et al. 2017). The first two PCA axes, which explained 63.5% of the total variance, were selected to calculate the hypervolumes. The boundaries of kernel density estimates were delineated by the probability threshold, using the 0.85-quantile value. Hypervolume contours were plotted to project niche spaces based on 5,000 random background points and using the alphahull (Pateiro-López and Rodríguez-Casal 2016) contour type and the alpha smoothing value set to 0.55. To evaluate statistically significant differences in mean climatic variables among species, we employed ANOVA analyses for each variable. Based on significant ANOVA results, we applied Tukey’s Honestly Significant Difference (HSD) test for post-hoc pairwise comparisons to identify which species pairs exhibited significant differences. All the analyses were performed in R Studio v. 6.2 (R Core Team 2024).

### 2.6. Species distribution modelling

To assess the current potential distribution of *Trebouxia decolorans* (A33), *T. solaris* (A35), and *T. tabarcae ad int.* (A48) —as the predominant symbiotic algal partners of *Xanthoria parietina*—species distribution modelling was conducted. The analysis included 19 bioclimatic variables (BIO1–BIO19) of the Worldclim database (Fick and Hijmans 2017), which represent temperature and precipitation patterns, along with altitude data.

Several modelling approaches were employed: generalized additive models (GAM) using the R package *gam* (Hastie 2024), generalized linear models (GLM) using the package *stats*, multivariate adaptative regression splines (MARS) using the package *earth* (Milborrow et al. 2024), maximum-entropy using MaxEnt v. 3.4.4 and the R package *maxnet* (Phillips 2021), random forest (RF) using the package *randomForest* (Liaw and Wiener 2002) and extreme gradient boosting training (XGBOOST) using the package *xgboost* (Chen et al. 2025). All the models were run within the package framework *biomod2* (Guéguen et al. 2025). Initially, all environmental variables were included in the models, and then, a selection of variables was performed based on their contribution to the models and their correlation among them. The final models were constructed using a reduced set of variables specific to each species: BIO4, BIO8, BIO11 and BIO14 for *Trebouxia decolorans* (A33); BIO4, BIO11 and BIO18 for *T. solaris* (A35); and BIO2, BIO6 and BIO7 for *T. tabarcae ad int*. (A48). Finally, model performance was evaluated using the area under the receiver operating characteristic curve (ROC) and the true skill statistic (TSS). Individual models were combined into ensemble models using a weighted mean based on their ROC and TSS, as implemented in the *biomod2* package.

### 2.7. Summed binary presence-absence maps

To visualize areas with overlapping potential distributions of *Trebouxia decolorans* (A33), *T. solaris* (A35), and *T. tabarcae ad int.* (A48), binary presence-absence maps were generated based on the ensemble probability outputs. For each species, ensemble predictions were converted to binary maps using two probability thresholds: 0.5 (50%) and 0.7 (70%). Grid cells with values equal to or higher than the threshold were classified as suitable (presence = 1), while the rest were classified as unsuitable (absence = 0).

The resulting binary rasters were summed pixel-wise to obtain combined presence maps, where each cell value ranges from 0 (no species predicted) to 3 (all three species predicted to occur). These maps represent regions where the potential distribution of one, two, or all the three species may overlap under current climatic conditions. All spatial analyses were conducted in R using the *raster* (Hijmans 2025a), *terra* (Hijmans 2025b), and *biomod2* packages.

## 3. Results

### 3.1. Phylogeny of Xanthoria mycobionts

A total of 174 newly obtained mycobiont sequences belonged to the genus *Xanthoria* (Figure 2). Although the sampling effort targeted thalli of *Xanthoria parietina* s. str., 32 of the collected thalli were finally molecularly identified as species other than *X. parietina*. Phylogenetic trees obtained through ML and Bayesian inference had congruent topologies and were consistent with previously published phylogenies of the genus (Arup et al. 2013; Khodosovtsev et al. 2023; Tsurykau et al. 2020). The resulting phylogeny, which included our newly generated *Xanthoria* sequences along with those obtained from the GenBank, revealed eight well-supported monophyletic clades. These corresponded to the seven previously recognized species —*X. aureola* s. lat., *X. ’ectaneoides’*, *X. mediterranea*, *X. monofoliosa*, *X. parietina*, *X. stiligera*, and *X. tendraensis*— and one novel lineage that we designated here as *Xanthoria* sp. ‘hydra’ (see below).Sequences of our samples were distributed across the clades of *X. parietina*, *X. aureola* s. lat., *X. monofoliosa*, and the novel *Xanthoria* sp. ‘hydra’. The monophyletic clade identified as *X. parietina* contained 142 of the 174 sequences from thalli that had been morphologically identified as *X. parietina*. The *X. monofoliosa* clade included 14 sequences blasting with ≥ 99% similarity to the reference sequences EU681293, EU681294 and JN984136. Seven new sequences formed their own distinct clade within the one encompassing *Xanthoria aureola* s. lat. sequences, which further included sequences of *X. calcicola* and *X. ectaneoides*. The newly identified clade *Xanthoria* sp. ‘hydra’, grouped 11 newly obtained sequences together with the GenBank sequence with accession number AJ320141, which corresponded to the specimen *Xanthoria* sp. SH87, collected in the Greek island of Hydra (Scherrer and Honegger 2003).

**FIGURE 2.**
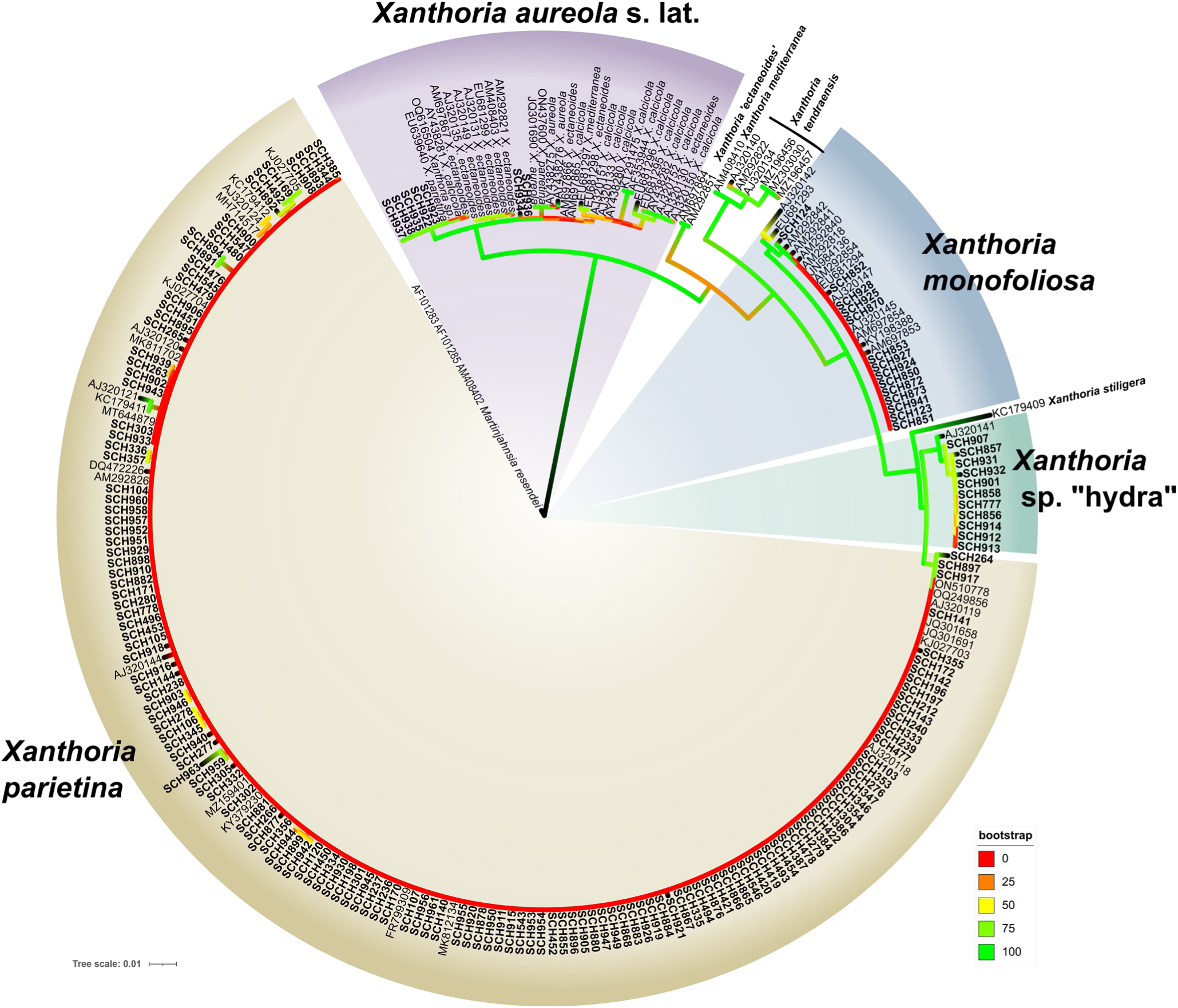
Phylogram generated using RAxML based on nrITS sequence data, showing the phylogenetic relationships among *Xanthoria* spp. mycobionts. Bootstrap support is represented by branches with different colors (see legend). GenBank accession numbers or sample codes are provided for each sequence/sample; additional details of the specimens are reported in Table S1.

### 3.2. Phylogenetic analysis of the *Trebouxia* phycobionts

All the phycobionts analysed in this study, as well as phycobiont sequences from *Xanthoria* lichens retrieved from the GenBank, were identified as species-level lineages within *Trebouxia* clade A (Figure 3). The topologies of both ML and Bayesian phylogenetic inferences were consistent with one another, and congruent with the most recent phylogenetic reconstructions of *Trebouxia* clade A (Chiva et al. 2025; Muggia et al. 2020).

All but three phycobiont sequences were grouped into nine different species-level lineages within *Trebouxia* clade A in *Xanthoria* spp. collected in the Mediterranean IbP and ItP. Eight of them corresponded to *Trebouxia decolorans* (A33), *T. solaris* (A35), *T. lynniae* (A39), *T. maresiae* (A46), *Trebouxia* sp. A13 *T. tabarcae ad int.* (A48), *Trebouxia* sp. A50, and *Trebouxia* sp. A66, while the ninth corresponded to a novel species-level lineage, here named *Trebouxia* sp. A56. Sequences belonging to this lineage did not correspond to any previously identified *Trebouxia* lineage at the species level and included two sequences from ItP, and one *Trebouxia* sequence from a *X. parietina* sample collected in the United Kingdom (GenBank accession no. ON453681).

In addition to this ninth *Trebouxia* spp., three Single-Occurrence Sequences (SOS) were reported (SCH276 as SOS1, SCH357 as SOS2, and SCH476 as SOS3). These sequences did not match any entry currently available in the GenBank, and form independent branches in the algal phylogeny (Figure 3). Although they belonged to the genus *Trebouxia*, SOS were not considered species-level lineages due to their singleton status, and they are therefore pending further detection in future studies before being formally recognized as taxonomic units. Nevertheless, they were included in the interaction network analyses (see below).

The most abundant phycobiont was *T. decolorans* (A33), whose clade allocated 70 sequences (47 from ItP and 23 from IbP). Nineteen sequences (nine from the ItP, ten from the IbP) were included in the monophyletic clade encompassing *T. solaris* (A35). Sixteen sequences (five from the ItP, 11 from the IbP) were included in *T. tabarcae ad int.* (A48). Eleven sequences (1 ItP, 10 IbP) were included in *Trebouxia* sp. A13, represented by the three paraphyletic species of *T. aggregata*, *T. arboricola* and *T. crenulata*. *Trebouxia maresiae* (A46) formed a highly supported clade and hosted four sequences from the IbP. The clades of *Trebouxia* sp. A50 (three sequences), *Trebouxia lynniae* (A39, three sequences) and *Trebouxia* sp. A66 (three sequences) were only found in three localities of the IbP.

Regarding climate preferences (see Table S1), 70 sequences of *T. decolorans* (A33) originated from areas with a Mediterranean climate on the Iberian (23 samples) and Italian (47 samples) peninsulas. Ten additional samples of *T. decolorans* were identified from areas outside the Mediterranean climate zone, specifically from France, Slovakia, and Norway—making it the only *Trebouxia* lineage in our dataset present in non-Mediterranean regions. These *Trebouxia decolorans* samples, together with the *T. tabarcae ad int.* (A48) samples, were found in Atlantic Europe. The other seven recognised *Trebouxia* species-lineages were only found in Mediterranean regions on both peninsulas.

Furthermore, 140 samples of *Xanthoria* spp. were collected as epiphytes, and only nine as saxicolous (see Table S1). The saxicolous ones associated with *Trebouxia* sp. A13 as the main phycobiont and corresponded to the following samples: one *X. parietina* collected on sandstones in Rende (loc. 11, ItP); one *X. parietina* and two *X. aureola* s. lat. collected on sandstones in Aín (loc. 35, IbP); and four thalli of *X. aureola* s. lat. collected on sandstones in Benicàssim (loc. 37, IbP). In the latter location, a fifth thallus contained *T. tabarcae ad int.* (A48) as the phycobiont.

**FIGURE 3.**
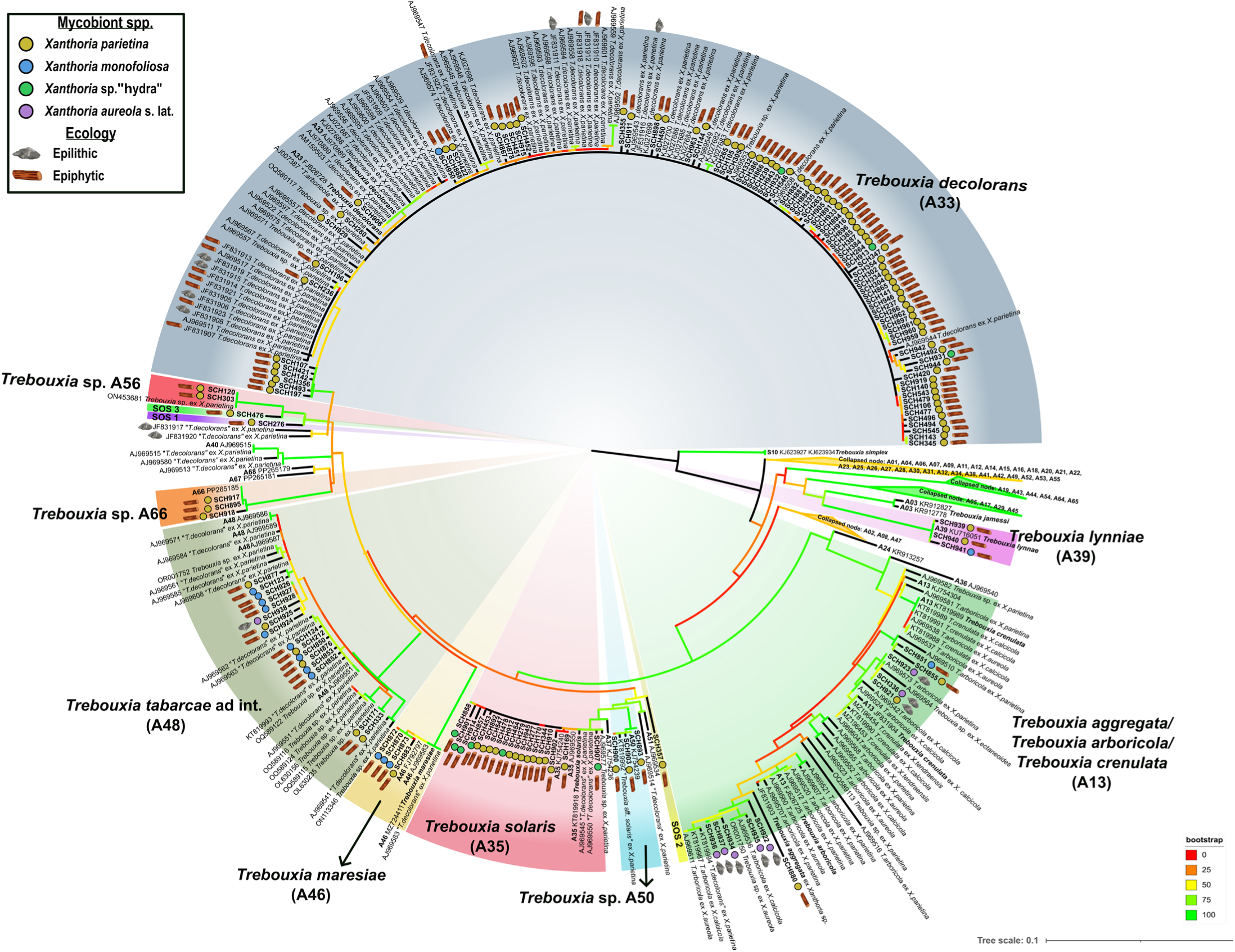
Phylogram generated using RAxML based on nrITS sequence data, representing the phylogenetic relationships between *Trebouxia* phycobionts of the analysed *Xanthoria* species. Bootstrap support is represented by branches with different colors (see legend). The substrate and the detected mycobiont lineages are indicated for each of the newly obtained sequences. Available information on the mycobiont lineage and substrate of the sequences retrieved from the GenBank was included (Figure 2; Table S2). GenBank accession numbers or sample codes are provided; additional details on specimens are reported in Table S1.

### 3.3. Interaction networks

Bipartite network analysis (Figure 4) highlighted that *Xanthoria parietina* was found in association with all nine identified *Trebouxia* species-level lineages, as well as the three SOS identified in the phylogenetic analysis (Figure 3). Despite this broad symbiotic range, *X. parietina* exhibited a strong preference for *T. decolorans* (A33). Other *Xanthoria* species displayed different predominant associations: *X. aureola* s. lat. was exclusively associated with *Trebouxia* sp. A13 and *T. tabarcae ad int.* (A48); *X. monofoliosa* associated with *T. lynniae* (A39), *T. maresiae* (A46), *Trebouxia* sp. A13 and *T. tabarcae ad int.* (A48); *Xanthoria* sp. ‘hydra’ associated with *T. decolorans* (A33) and *T. solaris* (A35). However, these results should be taken with caution, due to the lower sample size for *Xanthoria* species other than *X. parietina*.

**FIGURE 4.**
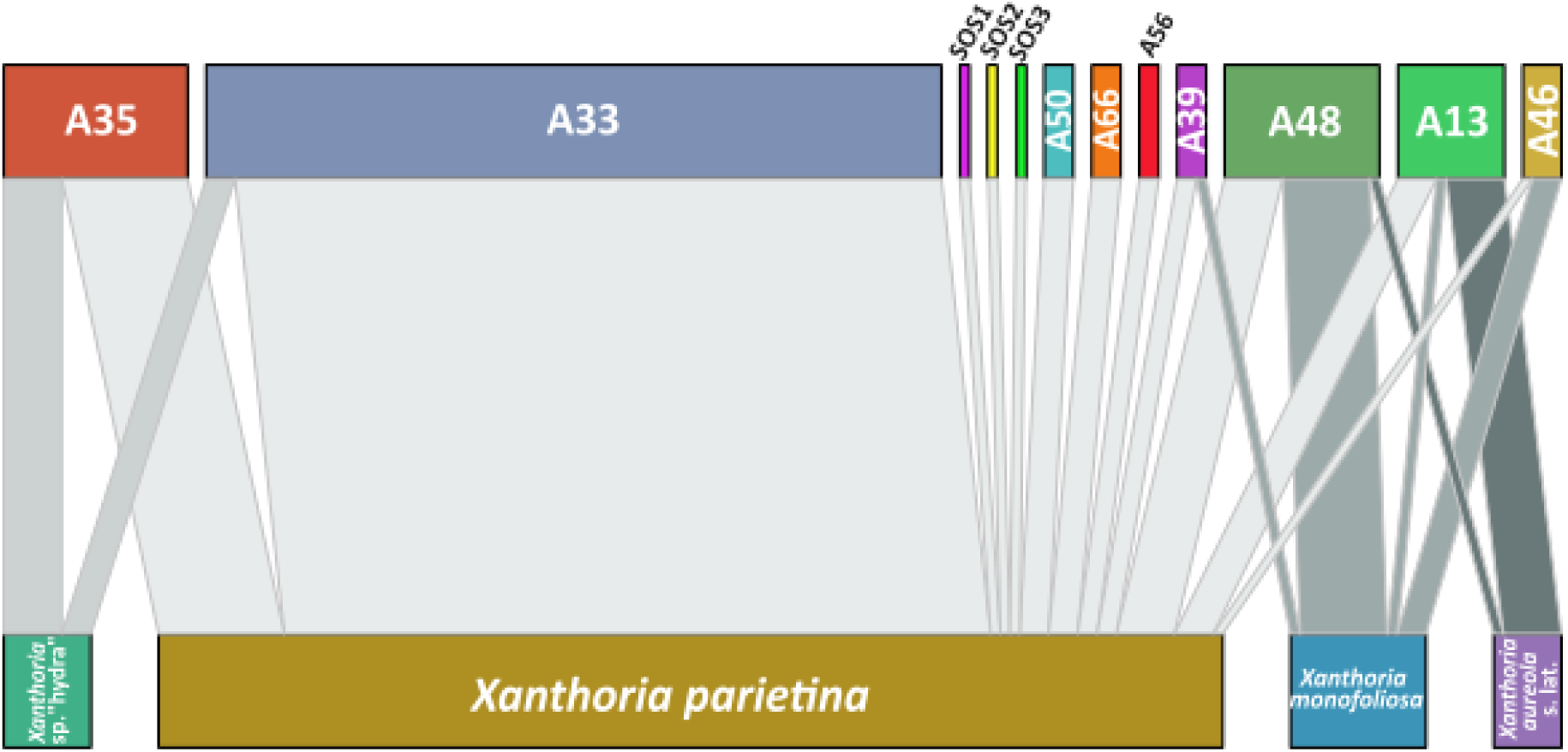
Interaction networks between the identified *Xanthoria* species and their *Trebouxia* phycobionts. *Trebouxia* taxa are reported with the numerical code of the corresponding species-level lineage in clade A. Single-Occurrence Sequences (SOS) are also included, although not considered species-level lineages. The width of the connections is proportional to the number of specimens representing the association.

The tripartite network (Figure S1) illustrates the associations between *Xanthoria* mycobionts and *Trebouxia* photobionts, and their geographic origin. *Xanthoria parietina* and *X. monofoliosa* were found in both regions, while *Xanthoria* sp. ‘hydra’ and *X. aureola* s. lat. were exclusively recorded in the IbP.

Among the photobionts, *T. decolorans* (A33) was the most commonly associated species in *X. parietina* thalli from the IbP and the ItP. Other lineages such as *T. solaris* (A35), *Trebouxia* sp. A13, and *T. tabarcae ad int.* (A48) were also present in both areas. In contrast, some *Trebouxia* lineages showed regional specificity: *Trebouxia* sp. A56 was found only in the ItP, whereas *T. maresiae* (A46), *T. lynniae* (A39), *Trebouxia* sp. A50, and *Trebouxia* sp. A66 were exclusively recorded in the IbP. Additionally, all SOS were recovered solely from the IbP. Interestingly, either the common *T. decolorans,* as well as *T. tabarcae ad int.* (A48), were also found in European Atlantic samples.

### 3.4. Climatic niche differentiation and distribution of Xanthoria parietina phycobionts

Ecological differentiation was analyzed for the three most abundant *Trebouxia* species associated with *X. parietina*, i.e., *Trebouxia decolorans* (A33), *T. solaris* (A35) and *T. tabarcae ad int.* (A48). Non-metric multidimensional scaling (NMDS) based on Bray-Curtis dissimilarities (Figure 5a) revealed distinct clustering patterns among the three lineages, with *T. solaris* (A35) and *T. tabarcae ad int.* (A48) showing closer spatial proximity between points. In contrast, *T. decolorans* (A33) exhibited broader distribution and ecological amplitude. Although the overlap between *T. solaris* (A35) and *T. tabarcae ad int.* (A48) was limited, a notable overlap in the hyperdimensional climate space was observed between *T. decolorans* (A33) and both *T. solaris* (A35) and *T. tabarcae ad int.* (A48).

The PCA of 19 WorldClim bioclimatic variables, along with altitude and distance to the sea, showed that the first principal components (PC) captured 63.2% of total variance. PC1 (39.9%) primarily represented thermal variation, while PC2 (23.3%) captured precipitation gradients. The resulting niche hypervolumes confirmed significant separation among the three species (e.g., BIO2: p = 6.21 × 10⁻⁵; BIO4: p = 0.0106; Figure S2).

*Trebouxia decolorans* (A33) occupied a niche centered around average temperature and precipitation values, with high tolerance for thermal variability (high BIO7), consistent with its widespread occurrence. *Trebouxia solaris* (A35) inhabited environments with relatively stable, moderate temperatures (indicated by low BIO7), but it had a preference for regions that receive higher precipitation, especially during typically dry periods (BIO14, BIO17, BIO18). The negative influence of BIO6 and BIO9 implied that this species may struggle in environments with very low temperatures during the coldest month or in extremely dry conditions.

*Trebouxia tabarcae ad int.* (A48) inhabited stable, warm environments with constant temperatures throughout the year (Figure 5b), as indicated by the negative influence of BIO4 and BIO7 and the high values for various temperature-related variables. This lineage was associated with high values in precipitation-related variables, particularly during both the coldest and warmest periods of the year.

**FIGURE 5.**
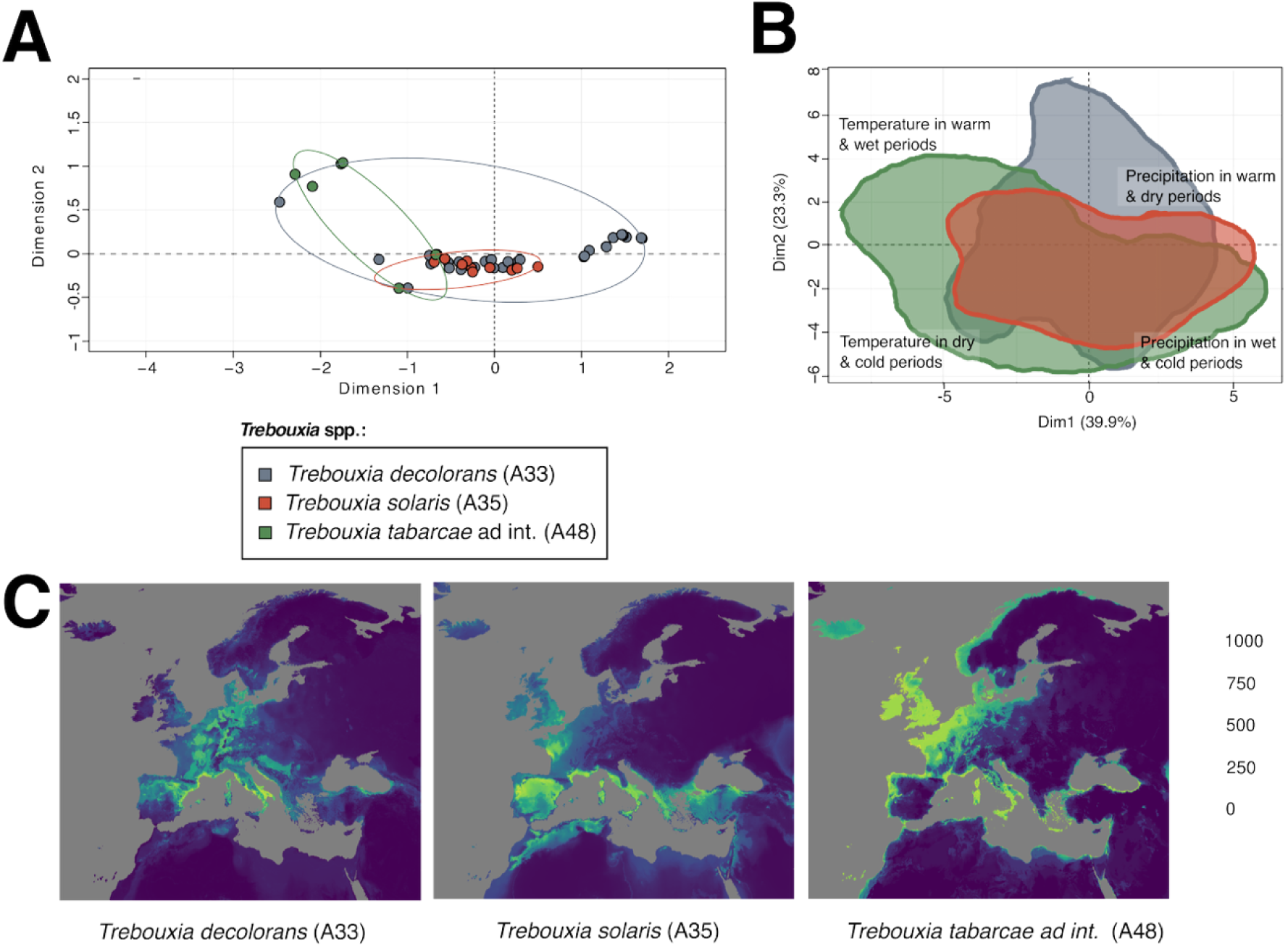
(A) Non-metric multidimensional scaling (NMDS) ordination plot of Bray-Curtis dissimilarities for *Trebouxia decolorans* (A33, blue), *T. solaris* (A35, red) and *T. tabarcae ad int*. (A48, green), present in thalli of *Xanthoria parietina*. (B) Climatic niche hypervolumes for *Trebouxia decolorans* (A33, blue), *T. solaris* (A35, red) and *T. tabarcae ad int*. (A48, green) based on climatic Dim1-Dim2 axes (63.2% of total variation explained). (C) Consensus distribution model for *Trebouxia decolorans* (A33), *T. solaris* (A35) and *T. tabarcae ad int*. (A48) in Europe based on MAXENT, MAXNET, GAMs, GLMs, MARS, RF and XGBOOST methods with selection of bioclimatic variables Colors range from dark blue (low suitability) to yellow (high suitability) according to the predicted habitat suitability index.

To predict the potential geographic ranges of these three *Trebouxia* taxa as phycobionts of *Xanthoria parietina*, seven modelling approaches (MAXENT, MAXNET, GAMs, GLMs, MARS, RF and XGBOOST) were employed to construct a consensus model based on the selected bioclimatic variables for *Trebouxia decolorans* (A33), *T. solaris* (A35), and *T. tabarcae ad int.* (A48) (see Figure S3 for variable selection). The receiver operating characteristic (ROC) and True Skill Statistic (TSS) values for each method are provided in Figure S4, with all models achieving high (> 0.80) ROC/TSS scores, reflecting strong predictive performance and model fit. Consistency among models generated by different methodologies was high (see Figure S5 detailing the resolution of each modelling method for all species). The consensus model of each species, depicted in Figure 5c, delineates areas of highest climatic suitability for *T. decolorans* (A33), *T. solaris* (A35), and *T. tabarcae ad int.* (A48) (see Figure S6 that outlines the relative contributions of each algorithm for each *Trebouxia* species). For *T. decolorans* (A33), the predicted distribution spanned broadly across Europe, covering extensive regions of Central Europe, the north-eastern IbP, and nearly the entire ItP, while avoiding high mountain ranges such as the Alps and the Pyrenees. In contrast, *T. solaris* (A35) showed a potentially wider distribution in Mediterranean areas, being particularly prevalent in the IbP, ItP, Mediterranean islands, Greece and Turkey (at approximately 40° N latitude) and the Atlas Mountains in Morocco. Furthermore, its distribution showed an Atlantic influence, with a notable presence in regions such as the English Channel and the Atlantic coasts in France and the IbP. It may also be frequent on all Mediterranean islands and along the coastline of the Mediterranean Basin. Additionally, there was a high likelihood of its presence across the United Kingdom and the North Atlantic coastal plain. The predicted distribution of *T. tabarcae ad int.* (A48) was slightly more restricted compared to the other taxa. Suitable climatic conditions may be concentrated in the western Mediterranean Basin, particularly in southern Spain and France, largely including coastal areas. Various isolated patches of high habitat favorability were also detected in northern Morocco and some zones of the central ItP. Unlike *T. decolorans* (A33) and *T. solaris* (A35), this species showed a minimally predicted presence in northern and central Europe, with limited potential of occurrence across high-altitude or Atlantic-influenced regions.

**FIGURE 6.**
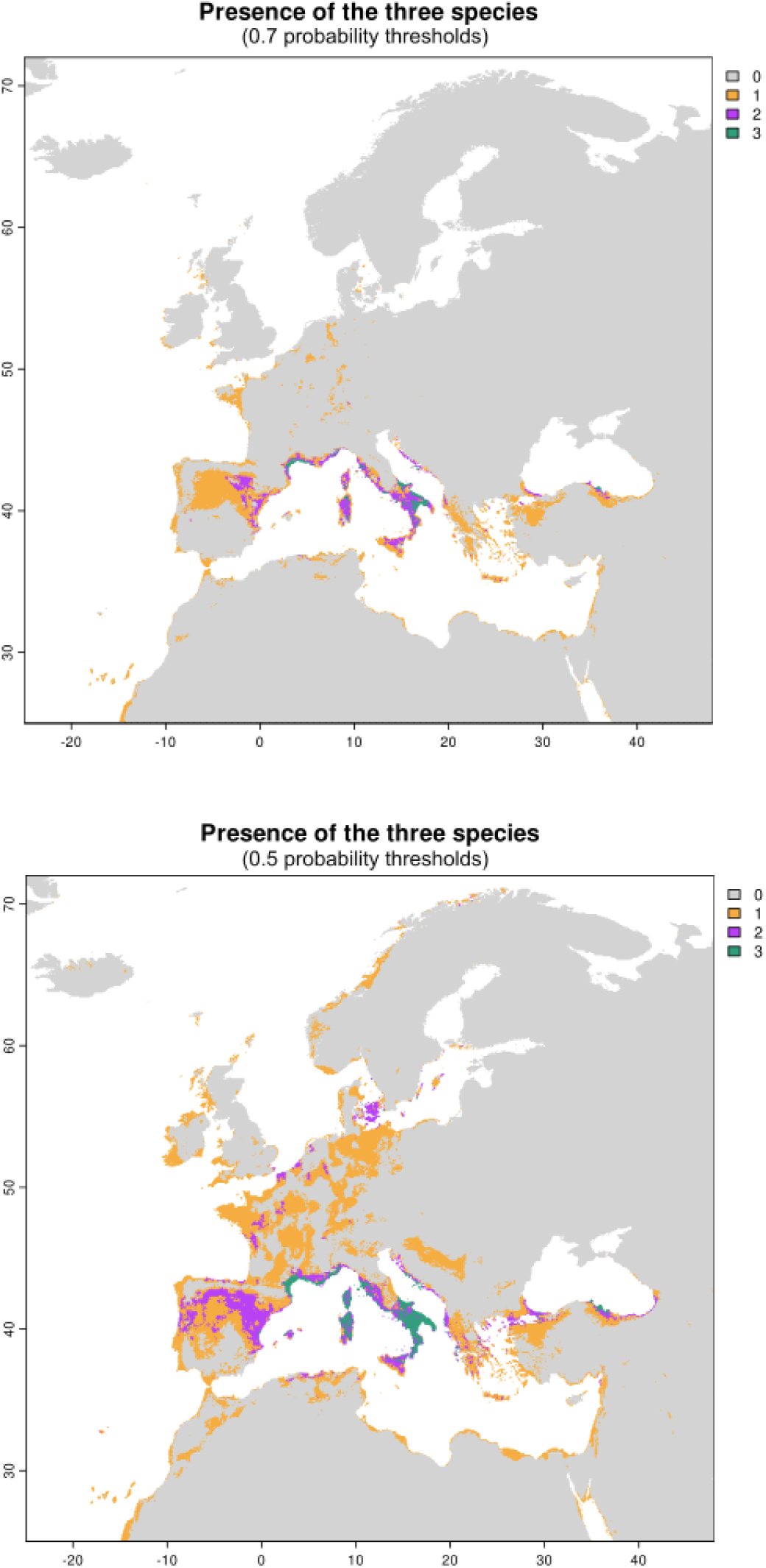
Predicted co-occurrence of the three studied Trebouxia species (T. solaris [A35], T. decolorans [A33], and T. tabarcae ad int. [A48]) in Europe and the Mediterranean Basin under two probability thresholds: 0.7 (top) and 0.5 (bottom). Colors indicate the number of species predicted to occur in each grid cell (0 = none, 1 = orange, 2 = purple, 3 = green).

The combined binary presence-absence maps (Figure 6) generated for the three selected *Trebouxia* species represented the predicted co-occurrence under current climatic conditions. At a 50% probability threshold, broader areas across Europe, particularly in temperate regions, showed predicted suitability for one or more *Trebouxia* species. The more conservative 70% threshold restricted their presence to more localized zones, highlighting areas with the highest environmental suitability and model agreement. Within these high-suitability areas, regions where all three species were predicted to co-occur (i.e., summed value = 3) included parts of northern Spain (especially the eastern Pyrenees), the coastal Mediterranean areas in France, the islands of Corsica and Sardinia, Tuscany and territories in southern Italy.

## 4. Discussion

Our results show that in the Mediterranean Basin the mycobiont *Xanthoria parietina* associates with a higher diversity of *Trebouxia* phycobionts than previously estimated. Although *X. parietina* is one of the most widespread and ecologically versatile lichen-forming fungi, capable of thriving on a wide variety of substrates and across diverse environmental conditions, the diversity of its microalgal partners has received limited attention (but see Beck and Mayr 2012; Dal Grande et al. 2014; Nyati et al. 2013, 2014; Wyczanska et al. 2023). Expanding knowledge of these associations is therefore crucial for understanding the ecological flexibility and the evolutionary dynamics of this almost cosmopolitan lichen species.

In this study, molecular data for accurate species identification proved essential to uncover both mycobiont diversity and their specific phycobiont associations. Molecular analyses showed that 32 of the 174 thalli initially identified as *X. parietina,* based on morphological features, were in fact different *Xanthoria* species. This finding underscores the limitations of morphology-based taxonomy within this genus, and highlights the importance of integrating molecular and morphological data for this group of lichen species (Kondratyuk et al. 2008). The considerable morphological and ecological variability within *X. parietina* has traditionally led to taxonomic confusion and the description of numerous forms and varieties (Hillmann 1920; Lindblom and Ekman 2005). The genus *Xanthoria* has undergone significant taxonomic restructuring in recent decades, with several species, such as *Rusavskia elegans,* reclassified based on molecular evidence (Kondratyuk and Kärnefelt 2003).

While molecular tools have greatly improved the accuracy of species identification, the widespread presence of mislabelled or poorly curated sequences in public genetic databases continues to generate biases and misinterpretations. For instance, in the case of *Xanthoria monofoliosa* S. Kondr. & Kärnefelt (Kondratyuk et al. 2008) —a species clearly delimited by the nrITS marker both by Fedorenko et al. (2009) and in the present analysis— only three sequences in the GenBank are correctly annotated. The remaining eleven sequences that form a well-supported monophyletic clade in our phylogeny are erroneously labelled as *X. parietina*, thereby distorting the abundance and distribution of either species.

In addition to *X. monofoliosa*, our study also revealed another associated taxon within the genus, identified as *Xanthoria* sp. “hydra”. This species has not yet been formally described and, until now, was represented by only a single sequence in the GenBank. With our new records, we highlight this taxon as potentially relevant and noteworthy for future description, as it occurs in several localities across the Iberian Peninsula.

### 4.1. Expanded phycobiont diversity in Xanthoria

Previous research had identified only four *Trebouxia* species —namely *T. decolorans*, *T. aggregata*, *T. arboricola*, and *T. crenulata*— as primary phycobionts of *X. parietina* (Nyati et al. 2014; Sanders and Masumoto 2021; Wyczanska et al. 2023). In contrast, our findings demonstrate that this mycobiont can associate with up to nine distinct *Trebouxia* species-level lineages, indicating a broader symbiotic range than previously recognized. According to Sanders and Masumoto (2021), *X. parietina* is exceptional among other lichens for its high phycobiont diversity, while other lichens, such as *Parmelia sulcata*, *Ramalina menziesii* or *Lecanora muralis*, despite exhibiting a relatively high photobiont diversity, present fewer and more restricted microalgae associations. To date, *X. parietina* appears to be the mycobiont with the highest number of known *Trebouxia* primary partners, highlighting its ecological and evolutionary versatility in forming lichen symbioses.

Comparable cases of clade-level microalgal diversity in *Trebouxia* have been linked to niche expansion in lichens (Blaha et al. 2006; Guzow-Krzemińska 2006; Medeiros et al. 2021). For example, *Cetraria aculeata* and *Pseudephebe* spp. shows a remarkable ability to colonize both polar and temperate habitats, and its broad distribution has been attributed to the ability of the mycobiont to associate with different *Trebouxia* lineages (Fernández-Mendoza et al. 2011; Garrido-Benavent et al. 2020). This suggests that photobiont flexibility can underpin the high colonization capacity of some lichens, allowing them to inhabit contrasting climatic regions. Such flexibility is also a determinant factor for other cosmopolitan lichens that colonize a wide variety of substrates — including tree bark, rocks and artificial materials, such as *Tephromela atra* or *Protoparmeliopsis muralis* (Kantnerová and Škaloud 2025; Muggia et al 2008, 2013, 2014a)—thriving from seashores to high mountains and in urban and polluted areas. This mechanism could also account for the wide distribution of *X. parietina*, whose ability to associate with different *Trebouxia* lineages may underlie its ecological success in both natural and anthropogenic environments.

In this context, it is important to note how *X. parietina* manages its symbiotic associations despite its broad ecological success. Although *X. parietina* reproduces sexually and is therefore likely to acquire photobionts opportunistically during spore germination and thallus development (Pichler et al. 2023), all the *Trebouxia* lineages identified in association with this mycobiont species belong to *Trebouxia* clade A. This indicates a certain degree of clade-level specificity, despite the wide range of microalgal partners involved. Similar specificity towards *Trebouxia* clade A has also been reported in other lichens, both sexually (apotheciate, reproducing by meiospores) and asexually (sorediate, isidiate, dispersing by diaspores) reproducing species (Leavitt et al. 2015; Moya et al. 2020, 2021; Muggia et al. 2014b). Further studies emphasize the role of reproductive strategy in shaping photobiont specialization in lichens (Berlinches de Gea et al. 2024; Steinová et al. 2019). This research has shown that, in Mediterranean forest lichen communities, sexually reproducing lichens generally exhibit lower specialization than asexual thalli, indicating that the reproductive strategy plays a major role in modulating symbiotic interactions in this environment (Berlinches de Gea et al. 2024). This pattern highlights that although sexual reproduction is often associated with broader, more opportunistic photobiont acquisition, it does not necessarily lead to phylogenetically unrestricted symbioses. In this sense, *Xanthoria parietina* provides a clear example: despite reproducing sexually and exhibiting a high diversity of associated photobionts, all the identified microalgae belong to *Trebouxia* clade A, suggesting a balance between ecological flexibility and phylogenetic specificity.

The previously overlooked *Trebouxia* species diversity taxa within *Xanthoria*, and in particular the discovery of new species-level lineages, underscores the importance of detailed investigation of lichen photobionts. To date, research on lichen symbiosis has been strongly biased towards the mycobiont partner (Muggia et al. 2018), although an increasing number of studies are highlighting the breadth of unexplored diversity among phycobionts, even in geographic regions where lichen species diversity has been investigated in depth, or for very common cosmopolitan lichens. Among the nine *Trebouxia* lineages identified in this study, several species-level lineages remain pending formal description, i.e., *T. tabarcae ad int.* (A48), *Trebouxia* sp. A50 and *Trebouxia* sp. A66. Alternatively, the complexity of *Trebouxia* sp. A13, represented by the three paraphyletic species *T. aggregata*, *T. arboricola*, and *T. crenulata*, requires taxonomic revision. The existence of undescribed but recurrently detected *Trebouxia* lineages in different studies, underscores the difficulty of describing species within this genus, largely due to the challenge of isolating axenically unialgal cultures (Chiva et al. 2021; Chiva and Moya 2025).

### 4.2. *Ecology of the* Trebouxia *phycobionts from* Xanthoria parietina

Climatic niche modelling has been applied to understand the variable processes responsible for shaping climatic tolerance in *Trebouxia*, and provides a framework within which to better understand potential responses to climate change-associated perturbations (Nelsen at al. 2022). Here, to understand the environmental drivers shaping the distribution of the *Trebouxia* phycobionts associated with *X. parietina*, we conducted ecological niche modelling to characterize their climatic requirements and preferences. Few studies have addressed the distribution of lichen symbiotic microalgae using hypervolume-based approaches. For example, Rolshausen et al. (2018) analysed the distribution of phycobionts in *Lasallia pustulata*, focusing on *Trebouxia* clade S (*Trebouxia angustilobata*, *T. simplex*, and other undescribed lineages), and demonstrated that distinct phycobionts exhibit divergent climatic tolerances. They showed that photobiont turnover along temperature and precipitation gradients likely facilitates fungal niche expansion, enabling *L. pustulata* to exploit a broader climatic habitat. Similarly, Moya et al. (2024) analysed *Ramalina farinacea* along a Mediterranean-Boreal gradient, characterizing interactions with *T. jamesii* and *T. lynniae* in *Ramalina* lichens. Their findings revealed overlapping but distinct climatic niches, with distributional segregation along climatic gradients, interpreted as evidence of environmental filtering acting on photobiont composition, suggesting that mycobionts may exploit alternative compatible partners to persist under varying environmental regimes, thereby expanding their ecological niches. These types of analyses are particularly valuable as they contribute to a better understanding of the environmental factors that may influence the selection of a particular microalgal partner by the mycobiont, as well as the processes that potentially induce algal switching (Piercey-Normore and DePriest 2001). Such insights shed light on the capacity of algal symbionts to colonize diverse habitats and illustrate how the replacement of a photobiont by a more locally adapted strain can facilitate colonization of new territories and enhance survival under adverse environmental conditions or climatic changes.

Our niche hypervolume analyses revealed significant differences among the symbiotic microalgae of *X. parietina*. For instance, *Trebouxia decolorans* (A33) exhibited a generalist profile, thriving in environments ranging from moderate temperature and humidity to areas with high thermal contrasts. In contrast, *Trebouxia solaris* (A35) seems to favor habitats that avoid extreme cold and dryness, while *T. tabarcae ad int.* (A48) appears to avoid regions with extreme temperature variability or pronounced seasonality. Apart from these differences, the overlap observed between some of the niche hypervolumes highlights *X. parietina*’s ability to switch its primary phycobiont depending on environmental conditions. In fact, even within a single locality, multiple *Trebouxia* species can be found as the main phycobionts of *X. parietina* —a phenomenon known as algal switching (Piercey-Normore and DePriest 2001)— which may represent a potential mechanism for environmental adaptation and niche evolution in mutualistic systems (Rolshausen et al. 2018; Škvorová et al. 2022). In our case, this process appears to be particularly relevant in Mediterranean environments, where it may enable the lichen to adapt to a broad spectrum of ecological conditions, including fine-scale microhabitats (Beck et al. 2002; Dal Grande et al. 2018; Ellis et al. 2021; Muggia et al. 2014b).

To compare the bioclimatic differences between the Mediterranean and other regions, we analysed phycobionts from *X. parietina* thalli collected across various bioclimatic zones in the Italian and Iberian peninsulas, and we incorporated phycobiont data from other *Xanthoria* species available in the GenBank. Across all our sampling localities outside the Mediterranean regions, *T. decolorans* was the only phycobiont identified, except in one Atlantic locality in the northern Iberian Peninsula (Euskadi), where *T. tabarcae ad int.* (A48) was detected. This pattern suggests that the symbiotic microalgal diversity in *X. parietina* is lower in non-Mediterranean European regions compared to Mediterranean ones. Such a pattern could reflect either a higher physiological compatibility or selectivity for *T. decolorans* in these environments, or a lower availability of alternative algal partners during thallus establishment. Notably, *T. decolorans* has been frequently reported as the sole phycobiont in lichens from Central Europe (Beck and Mayr 2012; Dal Grande et al. 2014; Voytsekhovich and Beck 2016; Wyczanska et al. 2023). This observation aligns with a general biodiversity pattern: the Mediterranean region is recognized as a hotspot of diversity and endemism, with higher species richness than temperate and continental areas of Europe (Coll et al., 2010; Comes 2004). Indeed, studies on *Ramalina farinacea* have shown greater variability in its phycobiont associations in the Mediterranean region, where different haplotypes of *Trebouxia* coexist, than in temperate and boreal areas (Moya et al. 2024). Similarly, recent reviews highlight that microalgal diversity in Mediterranean environments is high yet still underexplored, with significant biotechnological potential (Cosenza et al. 2024).

The finding that *Trebouxia decolorans* (A33) functions as a generalist symbiont and that the mycobiont *X. parietina* itself is highly adaptable (Lorenz et al. 2024), seems to explain the remarkable ecological plasticity of this lichen symbiosis, able to thrive under contrasting environmental conditions, from coastal habitats to high mountains, from Mediterranean to boreal areas, and from urban to pristine settings. However, under certain climatic conditions, other microalgae can also establish symbioses with *X. parietina* mycobiont, as they can be even better adapted than *T. decolorans*. Additionally, the availability of different free-living microalgae in these environments may influence the recruitment of phycobionts by *X. parietina* during its development.

The consensus distribution model allows the spatial estimation of potential distributions based on selected bioclimatic variables. In our Europe-based analyses, *T. decolorans* (A33) appears broadly distributed throughout Central Europe, while *T. solaris* (A35) shows a strong presence in the Mediterranean region. Remarkably, *T. tabarcae ad int.* (A48) is highlighted as being strongly localized in coastal areas of the Mediterranean. The presence of this alga as a coastal specialist has previously been reported in lichens such as *Seirophora villosa* (Garrido-Benavent et al. 2022) and *X. parietina* (Nyati et al. 2013, 2014; Voytsekhovich and Beck 2016). Similarly, among the photobionts of *X. parietina*, two other species—*Trebouxia lynniae* and *T. maresiae*—also show coastal distribution. Comparable results have been observed in *Ramalina farinacea* (Barreno et al. 2022) and *S. villosa* (Garrido-Benavent et al. 2022). Although this cannot be generalized, there is evidence that some *Trebouxia* taxa show distributions linked to particular climatic features (Werth and Sork 2014), and here a similar observation can be extended to the coastal distribution of *Trebouxia lynniae* and *T. maresiae*.

Our analyses are based on all available, rigorously curated data, although biases in sampling effort across phycobiont lineages might introduce uncertainty in hypervolume estimates (Jarvis et al. 2019; Werth and Sork 2014). As more high-quality occurrence records and ecological metadata on *Xanthoria* phycobionts become available, these hypervolume models will be refined, enabling more accurate predictions of each symbiotic microalga’s ecological niche and potential geographic range (Ascanio et al. 2024; Blonder 2018). In this context, the distribution maps of the three *Trebouxia* species illustrate how the choice of probability threshold strongly influences predicted presence areas. At a conservative threshold (0.7), suitable habitats are restricted to fragmented regions around the Mediterranean Basin, reflecting high-confidence predictions with the species’ ecological requirements. By contrast, a more permissive threshold (0.5) expands potential ranges considerably into central and northern Europe, revealing broader —but more uncertain— suitable conditions. These differences highlight both the importance of threshold selection in ecological niche modelling and the need for cautious interpretation, particularly when inferring species interactions or forecasting responses to climate change, as observed in studies of *Trebouxia* niche divergence (Nelsen et al. 2022).

### 4.3. Substrate, species, and regional patterns in the Trebouxia–Xanthoria symbioses

Substrate effects are likewise unsettled. Early work suggested that epiphytic thalli of *X. parietina* often host *T. decolorans,* whereas saxicolous thalli favour *T. arboricola* (Nyati et al. 2013). Other surveys in Central Europe (Beck and Mayr 2012) and a global metatranscriptomic study (Tagirdzhanova et al. 2025a) reported *T. decolorans* on both substrates. Our Mediterranean data corroborate this latter pattern: *T. decolorans* occurs in thalli collected from bark and rock. Two explanations remain plausible: (i) substrate effects may emerge only under certain climatic conditions; or (ii) previous substrate related patterns may reflect misidentified thalli belonging to other *Xanthoria* species.

Indeed, our dataset reveals clear species-specific photobiont preferences within the genus. *X. parietina* associates mainly with *T. decolorans*, a pattern already documented in previous molecular studies that identified this lineage as the dominant photobiont in *X. parietina* across different geographical scales (Dal Grande et al. 2014; Nyati et al. 2013). In turn, the mycobiont *X. monofoliosa* partners with *T. tabarcae ad int.* (A48) and *T. maresiae*, *Xanthoria* sp. ‘hydra’ selects *T. solaris*, and *X. aureola* prefers *Trebouxia* sp. A13, consistent with the tendency observed in other *Xanthoria* species, which show affinity for particular *Trebouxia* lineages of clade A (Nyati et al. 2014). Thus, although *T. decolorans* is the most frequent photobiont overall, it is absent from some congeners, reflecting genus-level specificity already reported in the lichen genus *Parmelia*, where different species groups display distinct associations with particular *Trebouxia* clades (Moya et al. 2021; Ossowska et al. 2019). Similarly, in *Ramalina farinacea*, a fixed pair of phycobionts (*Trebouxia jamesii* and *T. lynniae*) is consistently maintained throughout its distribution range, underlining the stability of species-specific associations in other lichens (Moya et al. 2024). These examples reinforce the conclusion that the association between *Xanthoria* and its *Trebouxia* partners follows a pattern of mycobiont–photobiont specificity, comparable to that described in various lichen genera (Medeiros et al. 2024). Geographical patterns reinforce this picture: in the tripartite network analysis (fungi–photobionts–regions), we detected the four *Xanthoria* species in the IbP versus two in the ItP and, in parallel, eight *Trebouxia* lineages in the IbP versus five in the ItP. In both regions, we observed shared core photobionts within *Trebouxia* clade A: *T. decolorans* (A33), *Trebouxia tabarcae ad int.* (A48), *T. solaris* (A35), and *Trebouxia* sp. A13. The broad ecological amplitude of *T. decolorans* as a photobiont, including its association with *X. parietina*, is well documented (Dal Grande et al. 2014), which explains its presence in both regions and its role as a connector in the symbiotic network.

The greater diversity of secondary lineages in the IbP could reflect (not mutually exclusive): (i) higher regional availability of free-living microalgae (Jung et al. 2024); and/or (ii) intrinsic patterns of specificity or tolerance by the mycobiont, modulated by environmental and genetic factors, as shown in eco-phylogenetic studies (Kosecka et al. 2022; Poquita-Du et al. 2024).

Future work should explicitly address these possibilities. In particular, integrative approaches combining morphology, physiology, and multilocus/phylogenomic datasets would be highly valuable to disentangle the complex symbiotic ecology of *Xanthoria* and to refine the taxonomy of *Trebouxia* lineages (e.g., the recent genomes of *Trebouxia lynniae* (Gazquez et al. 2024) or *Trebouxia tabarcae ad int.* (A48; Tagirdzhanova et al. 2025b). Such approaches would allow us to go beyond presence/absence data and begin testing the drivers of partner selection across regions.

## Supporting information

Supplementary material Chiva Xanthoria

## Acknowledgements

This work was supported by PROMETEO Excellence in Research Program (PROMETEO/2021/005 and PROMETEO/2024/044, Generalitat Valenciana, Spain) and Grant PID2021-127087NB-100 funded by MICIU /AEI/10.13039/501100011033 (Ministry of Science, Innovation, and Universities, Spain) and a Next Generation EU, Contract “Margarita Salas” to S. Chiva, MS21-058 (MCIU). Microscopy analyses were carried out at the microscopy facilities of the SCSIE (Servei Central de Suport a la Investigació Experimental) of the Universitat de València, Burjassot-Paterna Campus.

## Data Accessibility Statement

Sequences produced in this study are deposited in the NCBI GenBank under the accession numbers PX354354 to PX354530 and PX354725 to PX354870.

## Author Contributions

S.C: research development and leadership, sampling, molecular analysis, figures, writing, and review. T.P.: ecological niche modelling analyses, figures, R and software, writing and reviewing. J. M.-P.: ecological niche modelling analyses, figures, writing and review. P.M.: writing and review. I.G.-B.: sampling, writing and review. E.B.: funding, writing, and review. L.M.: sampling, funding, writing, and review.

