## Supplementary material Chiva Xanthoria for "Symbiotic versatility in action: *Trebouxia* diversity expands the niche of the lichen *Xanthoria parietina*": Supplementary Figure S3.pdf

Box plot for wc2.1 _30s_bio_2

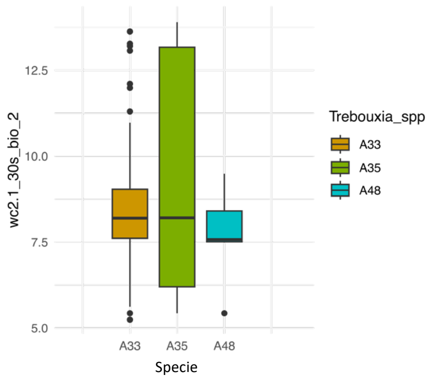


Box plot for wc2.1 _30s_bio_4

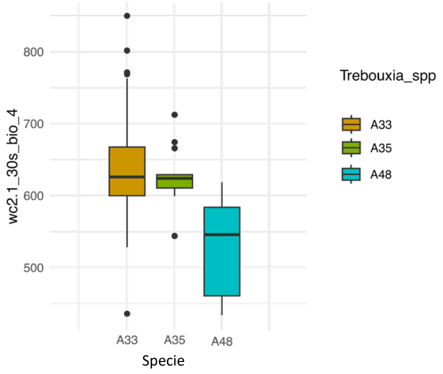


Box plot for wc2.1 _30s_bio_6

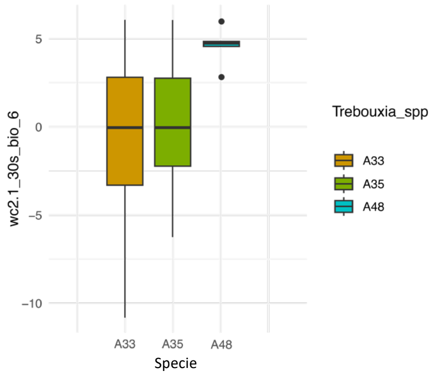


Box plot for wc2.1 _30s_bio_7

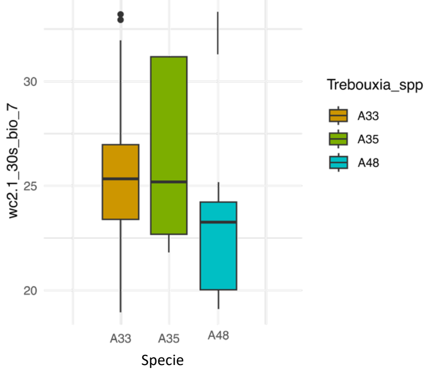


Box plot for wc2.1 _30s_bio_8

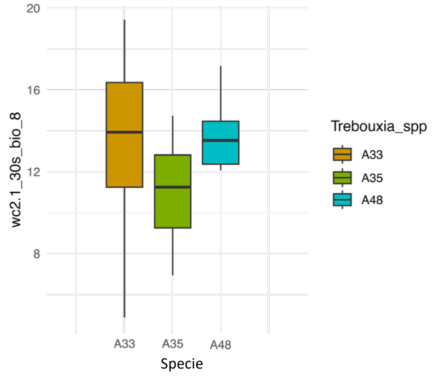


Box plot for wc2.1 _30s_bio_11

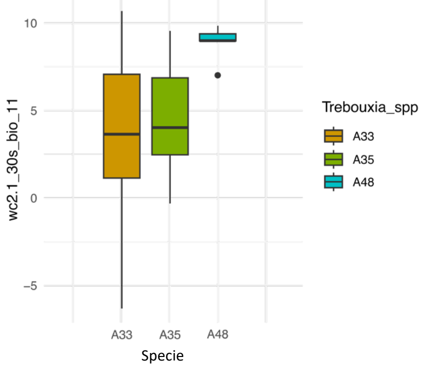


Box plot for wc2.1 _30s_bio_14

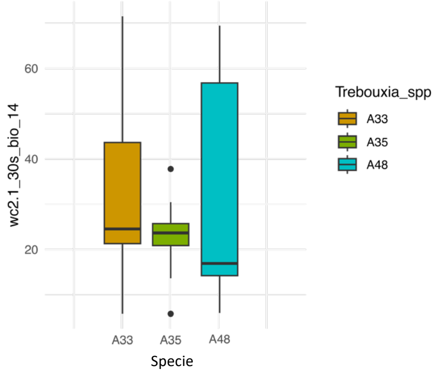


Box plot for wc2.1 _30s_bio_18

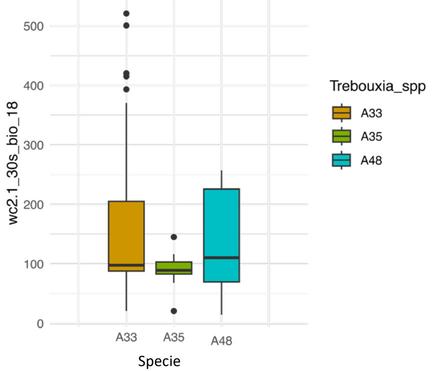
