## Supplementary figures and images for "Symbiotic versatility in action: *Trebouxia* diversity expands the niche of the lichen *Xanthoria parietina*"

### Supplementary Figure S1.tiff

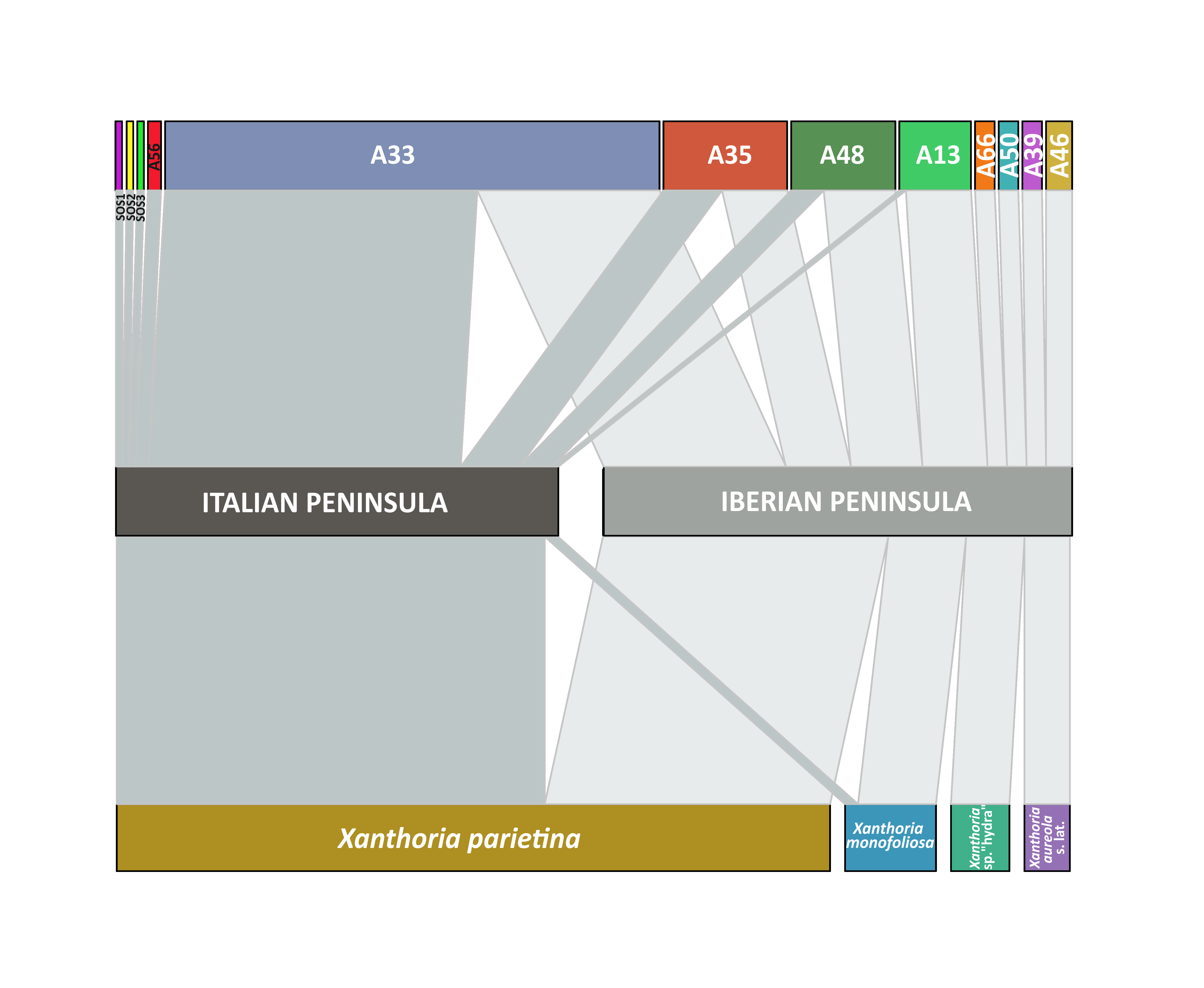

### Supplementary Figure S2.pdf

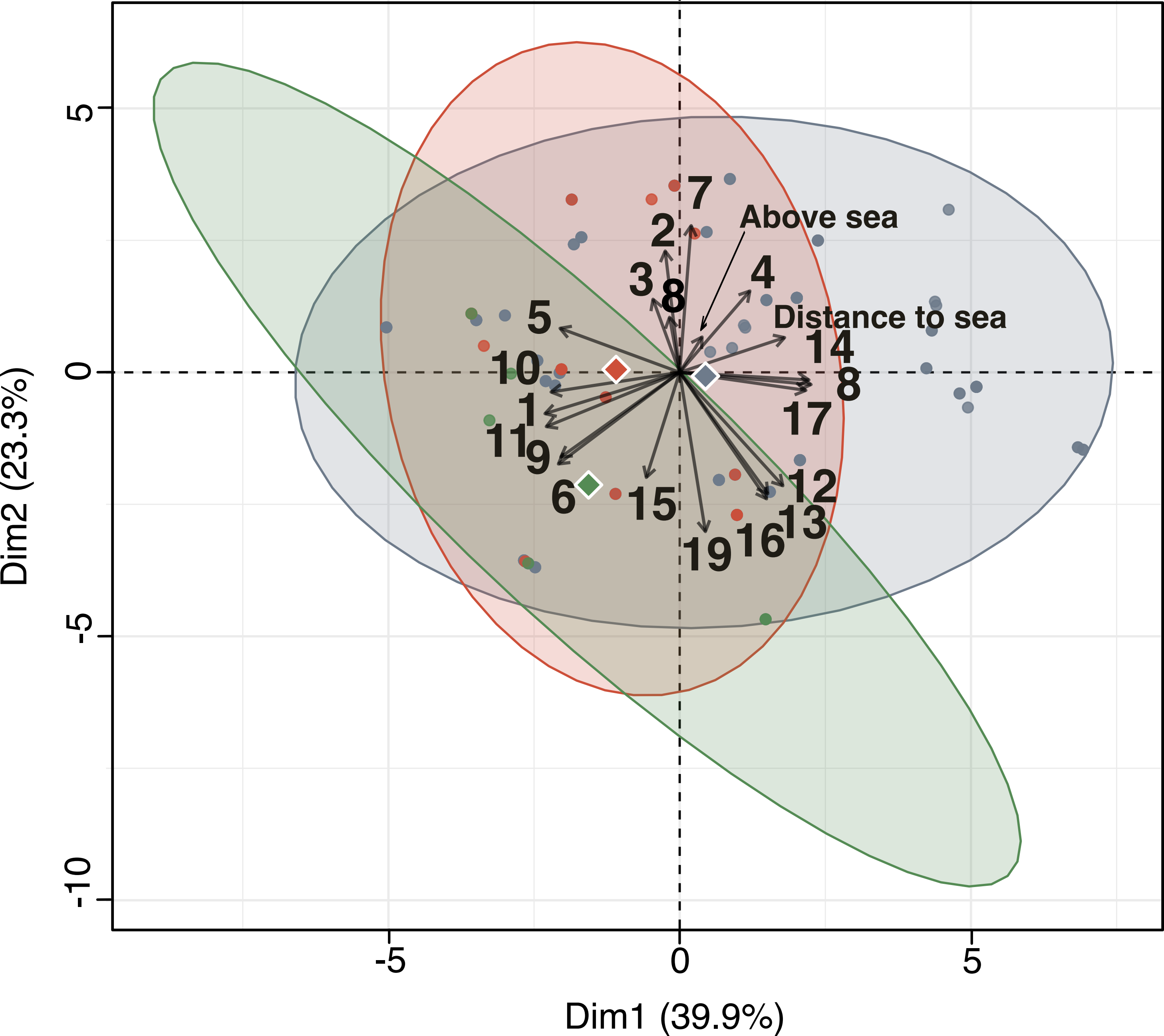

### Supplementary Figure S4.pdf

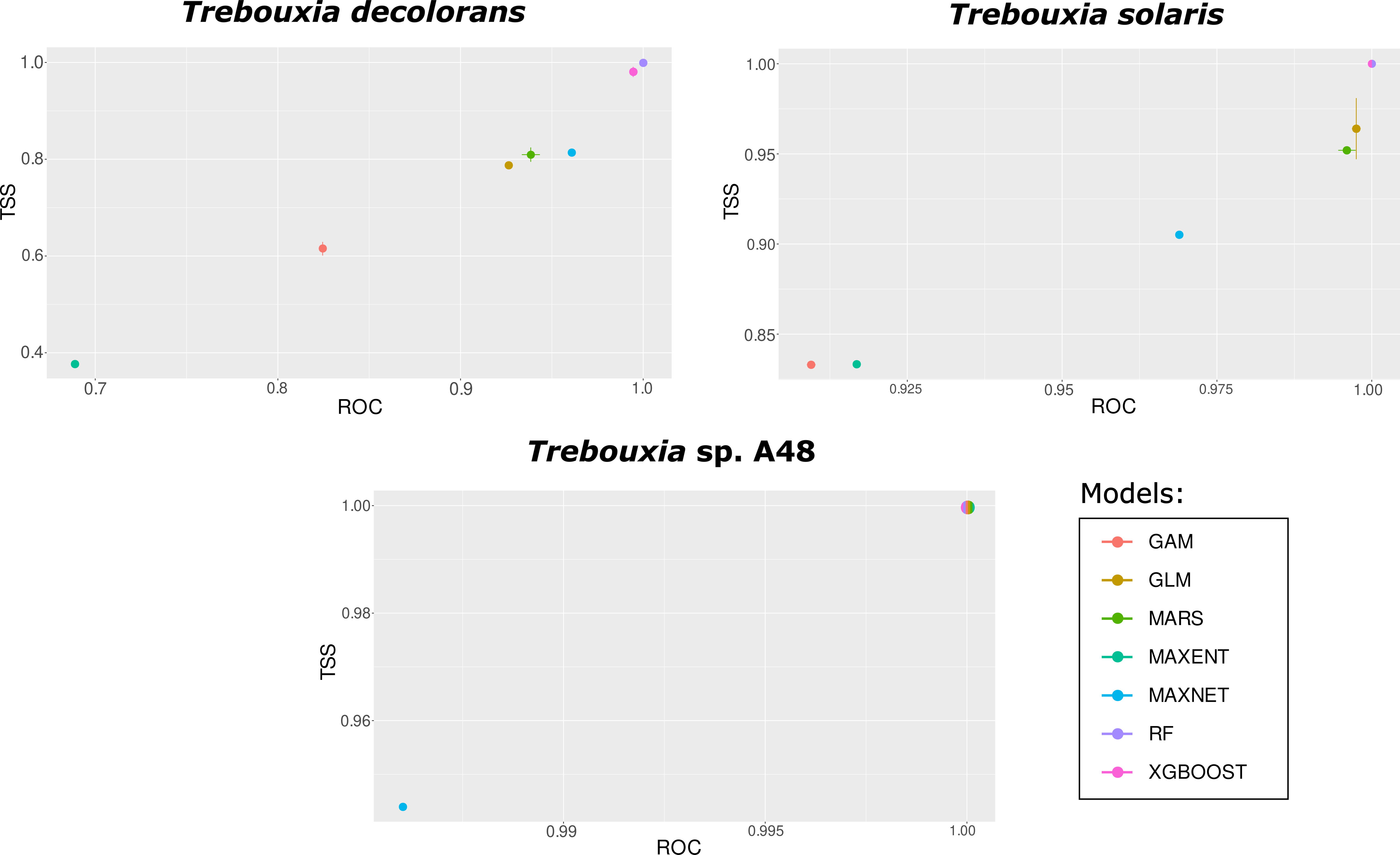

### Supplementary Figure S5.pdf

#### *Trebouxia decolorans* (A33)

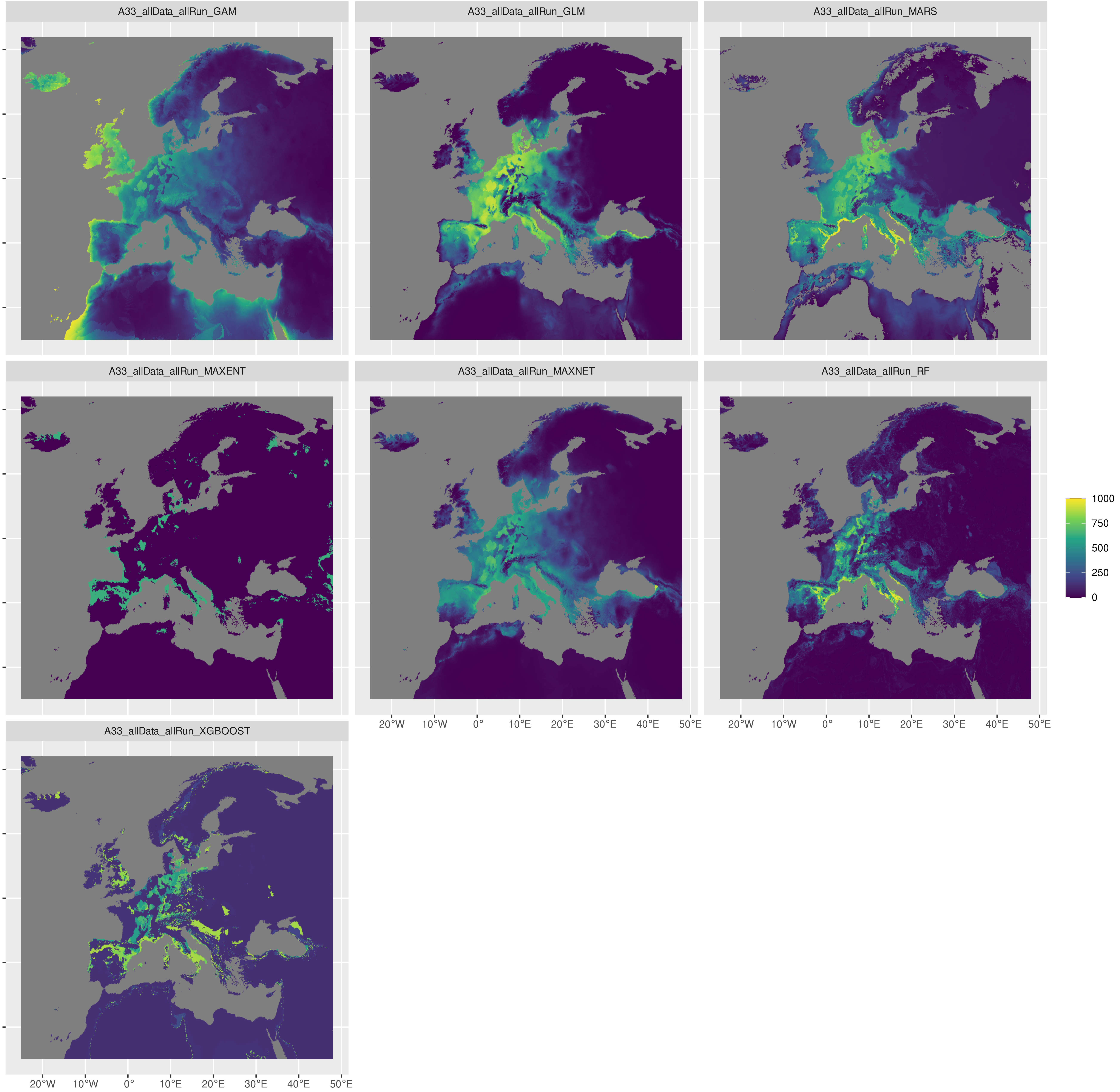

#### *Trebouxia solaris* (A35)

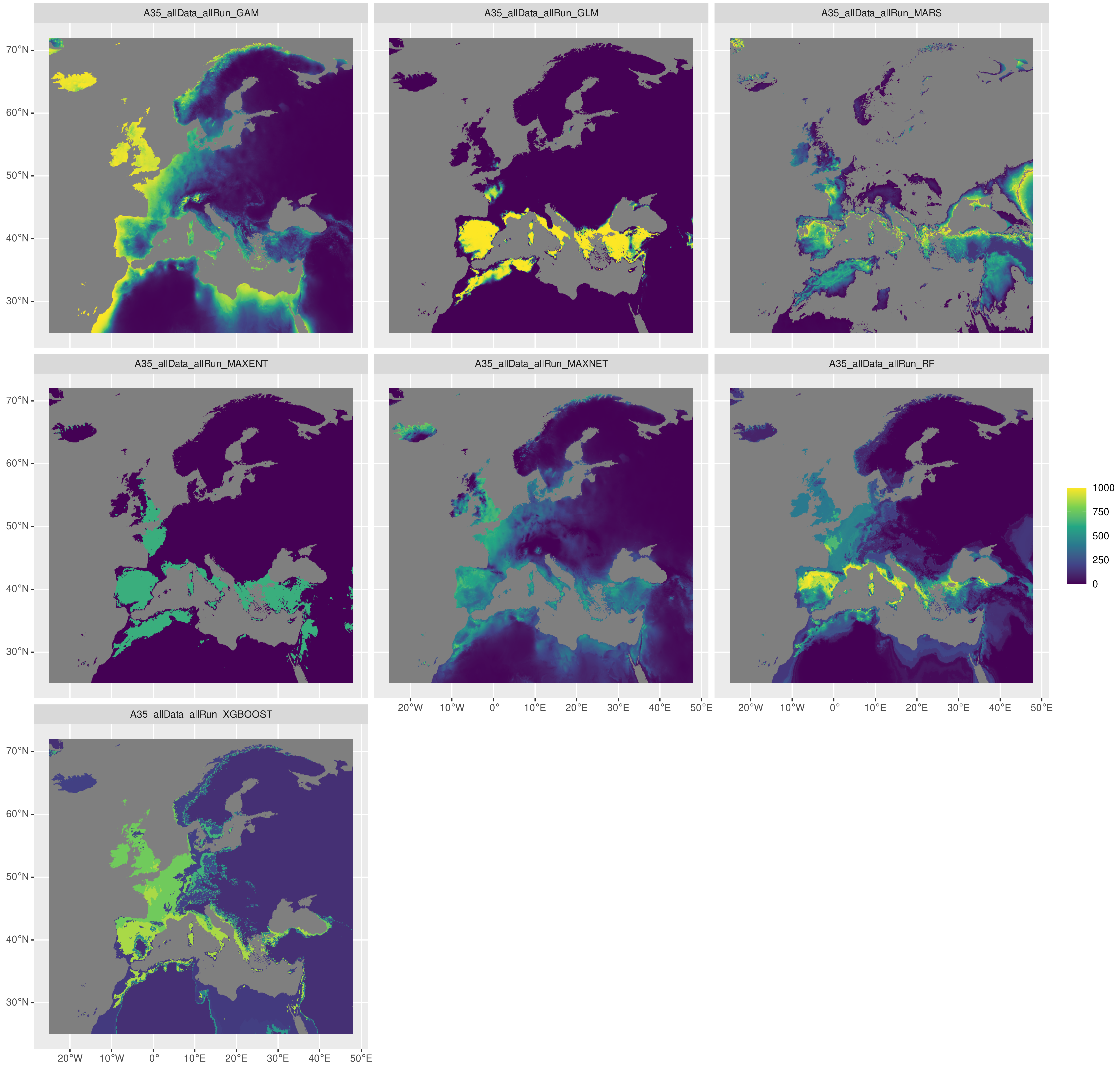

#### *Trebouxia* sp. A48

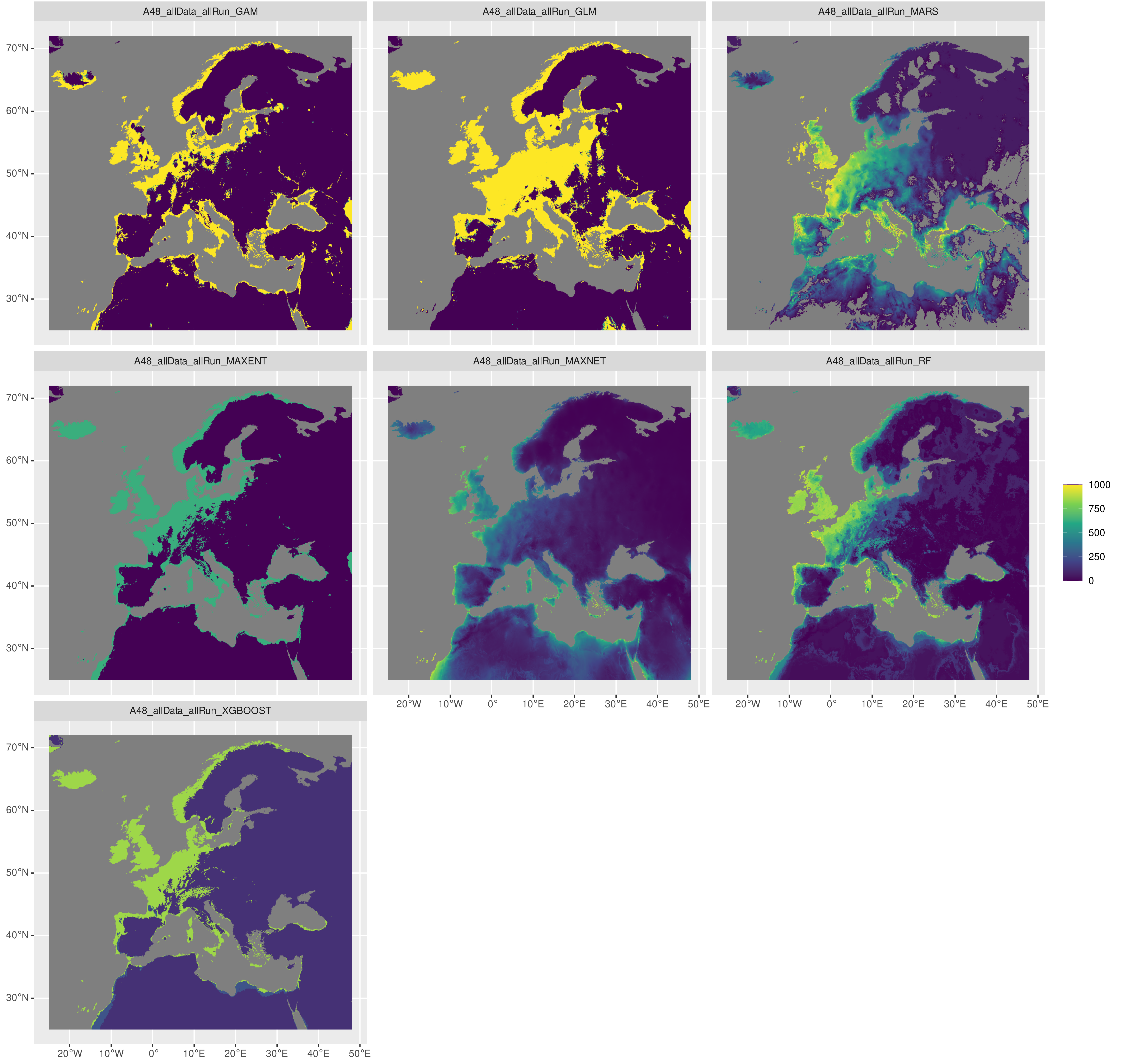

### Supplementary Figure S6.pdf

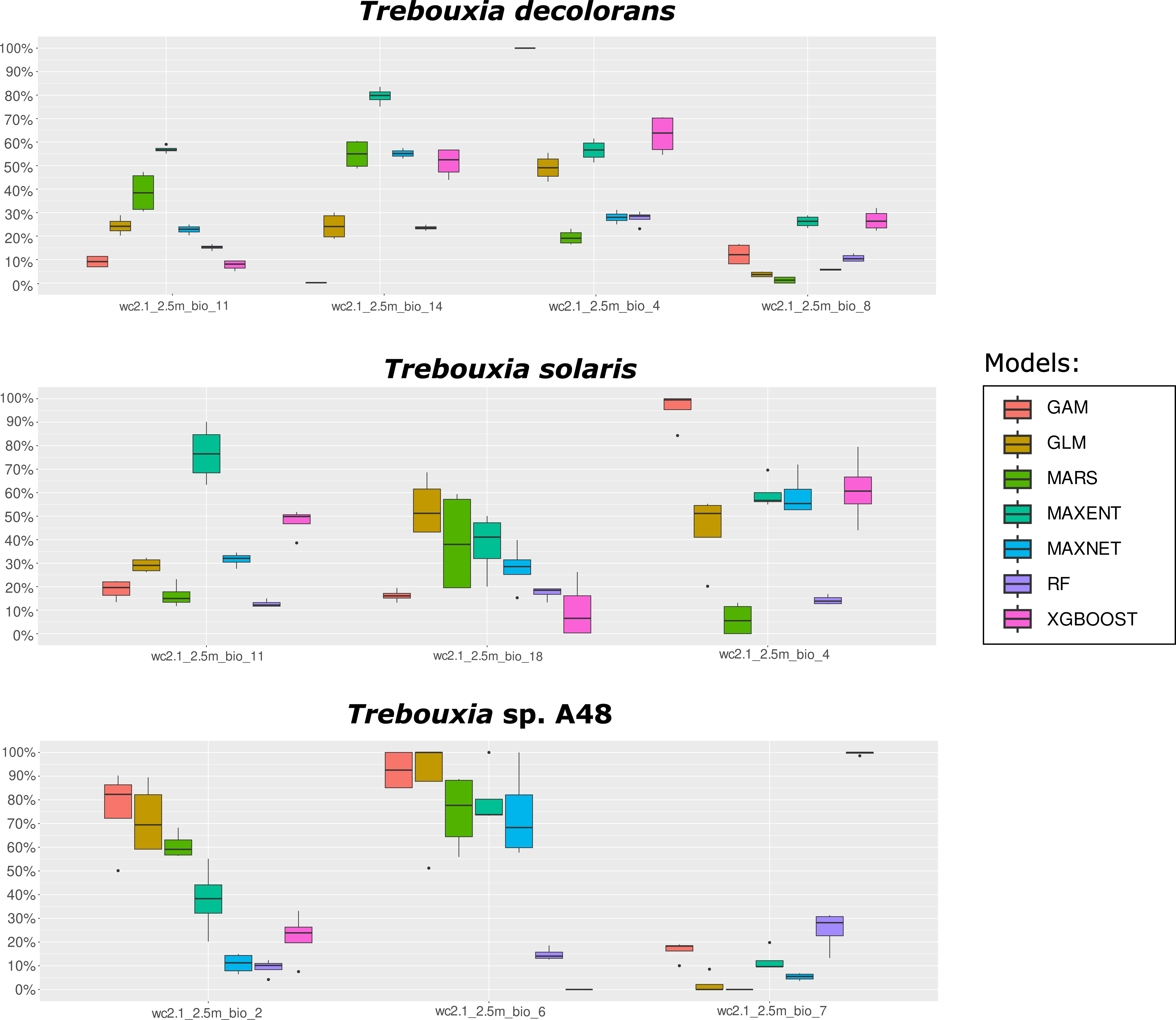
